# TRESK K^+^ Channel Activity Regulates Trigeminal Nociception and Headache

**DOI:** 10.1101/679530

**Authors:** Zhaohua Guo, Chang-Shen Qiu, Xinhua Jiang, Jintao Zhang, Fengxian Li, Qin Liu, Ajay Dhaka, Yu-Qing Cao

## Abstract

Although dominant-negative mutations of TRESK background K^+^ channel have been reported in migraine patients, whether TRESK activity controls the generation of trigeminal pain, especially headache, is not established. We found that loss of TRESK in all trigeminal ganglia (TG) neurons preferentially increased the intrinsic excitability of small-diameter TG nociceptors that express neuropeptide CGRP or TRPM8 channels. Surprisingly, loss of TRESK increased the number of TG neurons expressing TRPV1 channels. In dorsal root ganglia neurons, neither the persistent outward current nor the intrinsic excitability was affected by the loss of TRESK. Compared with wild-type controls, TRESK knockout mice exhibited more robust trigeminal pain, especially headache-like behaviors; but displayed normal body and visceral pain responses. Our findings indicate that endogenous TRESK activity is required for trigeminal pain regulation. A substantial reduction of TRESK activity may selectively affect the functions of TG nociceptors, thereby increasing the susceptibility to migraine headache in humans.

## Introduction

The TWIK-related spinal cord K^+^ (TRESK) channel belongs to the two-pore domain K^+^ (K_2P_) family of background K^+^ channels and is encoded by a single gene KCNK18 (Enyedi & Czirjak, 2015; Kang et al., 2004; Sano et al., 2003). Of all 15 mammalian K_2P_ channels, TRESK is the only one that is abundantly expressed in primary afferent neurons (PANs) in trigeminal ganglia (TG) and dorsal root ganglia (DRG) but at a negligible level in other tissues, suggesting that its main physiological function is to control somatosensation via regulating PAN excitability (Czirjak et al., 2004; Dobler et al., 2007; Kollert et al., 2015; Lafreniere et al., 2010; Yoo et al., 2009). Indeed, previous studies indicate that TRESK is one of the major background K^+^ channels in DRG neurons (Kang & Kim, 2006; Marsh et al., 2012). It is well established that endogenous TRESK activity regulates the excitability of DRG neurons and controls the transmission of body pain under both normal and chronic pain conditions. Both chronic nerve injury and tissue inflammation reduce the level of TRESK mRNA in rat DRG (Marsh et al., 2012; Tulleuda et al., 2011). Inhibition of TRESK and other K_2P_ channels by sanshool robustly increases the firing of DRG mechanoreceptors (Bautista et al., 2008; Lennertz et al., 2010). Reducing DRG TRESK activity elicits nocifensive behavior and increases mechanical sensitivity on the hindpaw (Tulleuda et al., 2011; Zhou et al., 2016). Over-expression of TRESK in rat DRG neurons inhibits neuropeptide release and attenuates neuropathic pain (Zhou et al., 2013; Zhou et al., 2012).

Much less is known about the function of TRESK channels in controlling trigeminal pain transmission, especially the trigeminovascular pathway subserving headache generation. Dominant-negative TRESK mutations are identified in patients with migraine but not with other chronic pain (Lafreniere et al., 2010), suggesting that genetic TRESK dysfunction differentially affects the generation of trigeminal pain and body pain. However, a recent study suggests that TRESK mutations increase TG neuronal excitability not through reducing TRESK current *per se*, but though inhibition of other K_2P_ channels TREK1/2 (Royal et al., 2018), calling into question the role of endogenous TRESK activity in controlling headache and other trigeminal pain. Many outstanding questions remain unanswered. Does endogenous TRESK activity regulate the excitability of dural afferent neurons, the PANs in the trigeminovascular pathway? Does TRESK dysfunction enhance trigeminal nociception, especially headache generation, at the behavioral level? In contrast to injury-induced reduction of TRESK activity in adults, can DRG neurons tolerate genetic TRESK dysfunction and maintain normal intrinsic excitability? If so, does this result in normal responses to stimuli that evoke body pain?

Here, we used TRESK knockout (KO) mice to investigate how endogenous TRESK activity regulates trigeminal nociception; whether genetic loss of TRESK differentially affects the generation of trigeminal and body pain. We found that ubiquitous loss of TRESK in all KO TG and DRG neurons preferentially increases the intrinsic excitability of small-diameter TG nociceptors that express calcitonin gene-related peptide (CGRP) or transient receptor potential channel melastatin 8 (TRPM8). Unexpectedly, the percentage of neurons expressing transient receptor potential cation channel subfamily V member 1 (TRPV1) channels, a sensor for a variety of noxious stimuli, is significantly increased in the TG but not DRG of KO mice. Compared with wild-type (WT) mice, TRESK KO mice exhibit more robust behavioral responses in various models of trigeminal pain, including headache. But their responses in body pain or visceral pain models are similar to those of WT mice. Our finding is the first demonstration that TG neurons are more vulnerable to TRESK dysfunction than DRG neurons; and genetic loss of TRESK preferentially enhances trigeminal pain.

## Results

### TRESK protein is ubiquitously expressed in PANs

TRESK KO mice (Figure 1A) were grossly normal. Progenies from heterozygous (HET) crossing had the expected Mendelian frequency. Both male and female KO mice gained weight normally and were fertile, with normal litter size and maternal behavior. First, we examined the distribution of TRESK protein in TG and DRG tissues. Previous studies using *in situ* hybridization and immunohistochemistry approaches indicated that TRESK is widely expressed in PANs (Dobler et al., 2007; Guo & Cao, 2014; Kollert et al., 2015; Lafreniere et al., 2010; Yoo et al., 2009); whereas single cell RNA-sequencing (RNA-seq) studies reported a more restricted TRESK mRNA expression in subpopulations of PANs (Chiu et al., 2014; Li et al., 2016; Usoskin et al., 2015). We stained TG and DRG sections from adult WT and KO mice with an antibody that recognizes mouse TRESK protein (Guo & Cao, 2014; Liu et al., 2013). TRESK-immunoreactivity (TRESK-ir) was completely absent in tissues from KO mice (Figure 1B), validating the specificity of the antibody. In TG and DRG sections from WT mice, we used an βIII tubulin antibody to label all neurons (Golden et al., 2010) and found that TRESK-ir overlapped with βIII tubulin signal in almost all neurons (Figure 1B), indicating that TRESK protein is ubiquitously expressed in mouse PANs but not in non-neuronal cells. It is possible that the level of TRESK mRNA is low in individual neurons, and single-cell RNA-seq may underestimate the number of neurons that express TRESK proteins.

**Figure 1.**
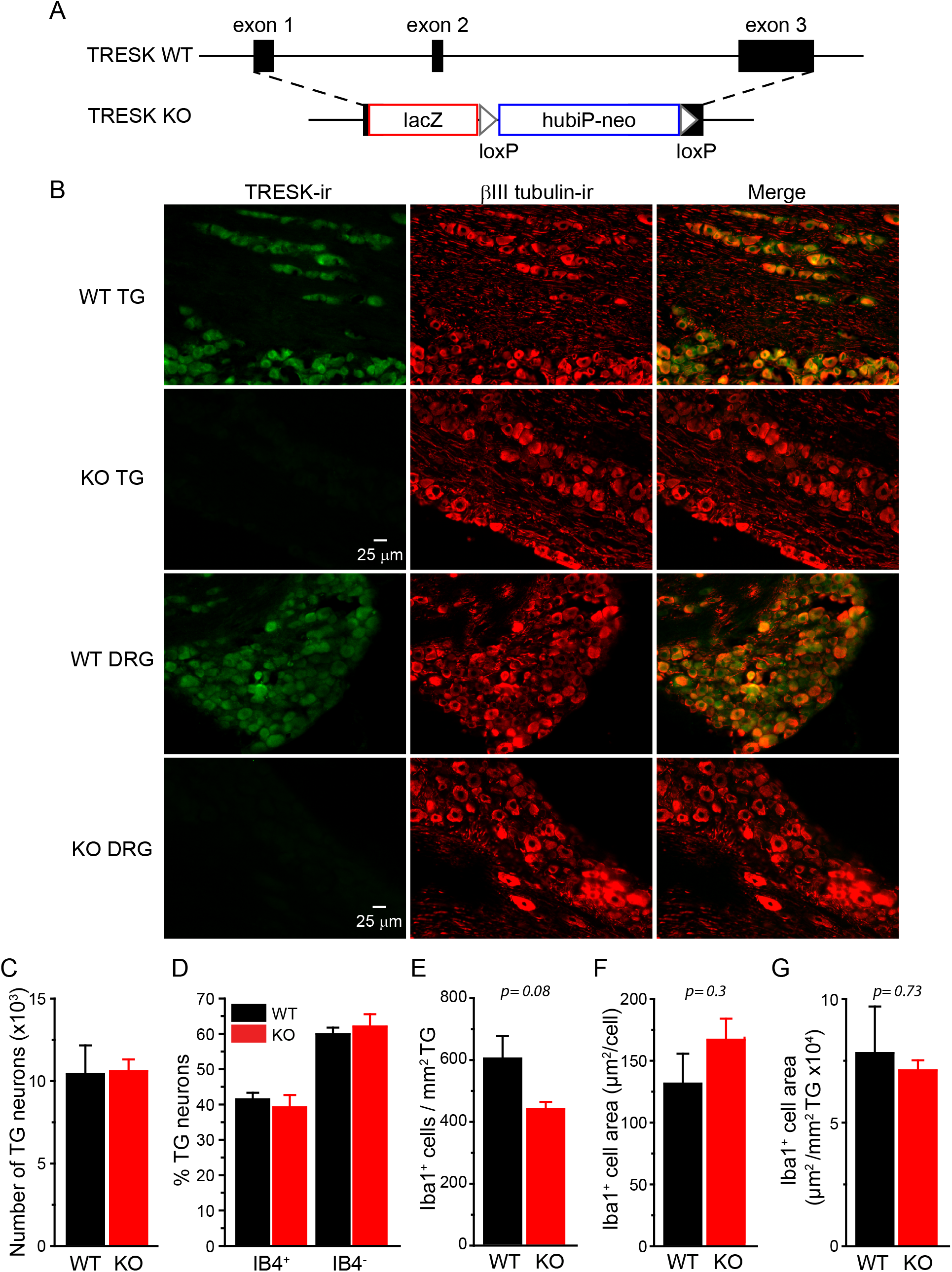
Functional TRESK channels are present in all TG and DRG neurons. (A) Schematics of WT mouse TRESK/Kcnk18 allele and the targeted allele. After homologous recombination, the lacZ-loxP-hubiP-neo-loxP cassette replaces most of the TRESK coding region. (B) Representative images of TG and DRG sections from adult WT and TRESK KO mice double stained with antibodies against TRESK and βIII tubulin. βIII tubulin-ir is present in all neurons. TRESK-ir is present in almost all neurons in the WT sections. There is no TRESK-ir in the KO sections. (C) Total number of TG neurons in WT and TRESK KO mice (n = 3 mice in each group). (D) The percentage of IB4^+^ and IB4^−^ neurons in WT and KO TG culture (1010 WT and 940 KO TG neurons from 10 separate primary cultures were counted). (E) The density of Iba1-positive (Iba1^+^) macrophages in WT and KO TG (n = 3 mice in each group). (F) The mean area of individual Iba1^+^ macrophages in WT and KO TG (same mice as in E). (G) The area of Iba1^+^ macrophages per mm^2^ TG in WT and KO mice (same mice as in D).

Next, we asked whether loss of TRESK affects the gross development of TG in mice. The total number of TG neurons containing βIII tubulin-immunoreactivity (βIII tubulin-ir) did not differ between adult WT and TRESK KO mice (Figure 1C), suggesting that TRESK is not required for the survival of PANs. The abundance of TG neurons that binds to the plant isolectin B4 (IB4) was also comparable between cultured TG neurons from adult WT and KO mice (Figure 1D).

In a mouse model of familial hemiplegic migraine type 1, the number of macrophages was significantly increased in TG, which may contribute to the headache pathophysiology (Franceschini et al., 2013). We stained macrophages in adult WT and KO TG with an antibody recognizing ionized calcium binding adaptor molecule 1 (Iba1). Both the density and the area of macrophages were similar between wild-type and KO TG (Figures 1E-G). We next focused on investigating how loss of TRESK affects the functions of PANs and the behavioral responses to noxious stimuli.

### Ubiquitous loss of TRESK currents in PANs preferentially increases the excitability of small-diameter TG nociceptors that do not bind to IB4 (IB4**^−^**)

In previous studies, we used the sensitivity to lamotrigine to dissect currents through TRESK channels (Guo & Cao, 2014; Guo et al., 2014; Liu et al., 2013). Bath application of 30 µM lamotrigine blocked 20-30% of persistent outward currents in every adult WT TG neurons that we recorded, regardless of soma size (Figures 2A-C, D, F). The same concentration of lamotrigine did not significantly reduce the outward currents in TG neurons from TRESK KO mice (Figures 2B-C, D, F), indicating that the majority of lamotrigine-sensitive current in TG neurons are mediate through TRESK channels under our recording conditions. The size of total persistent outward current was also significantly reduced in KO TG neurons (Figures 2E, G). TG neurons from HET mice showed intermediate levels of lamotrigine-sensitive TRESK current and total persistent outward current (Figures 2C-E).

**Figure 2.**
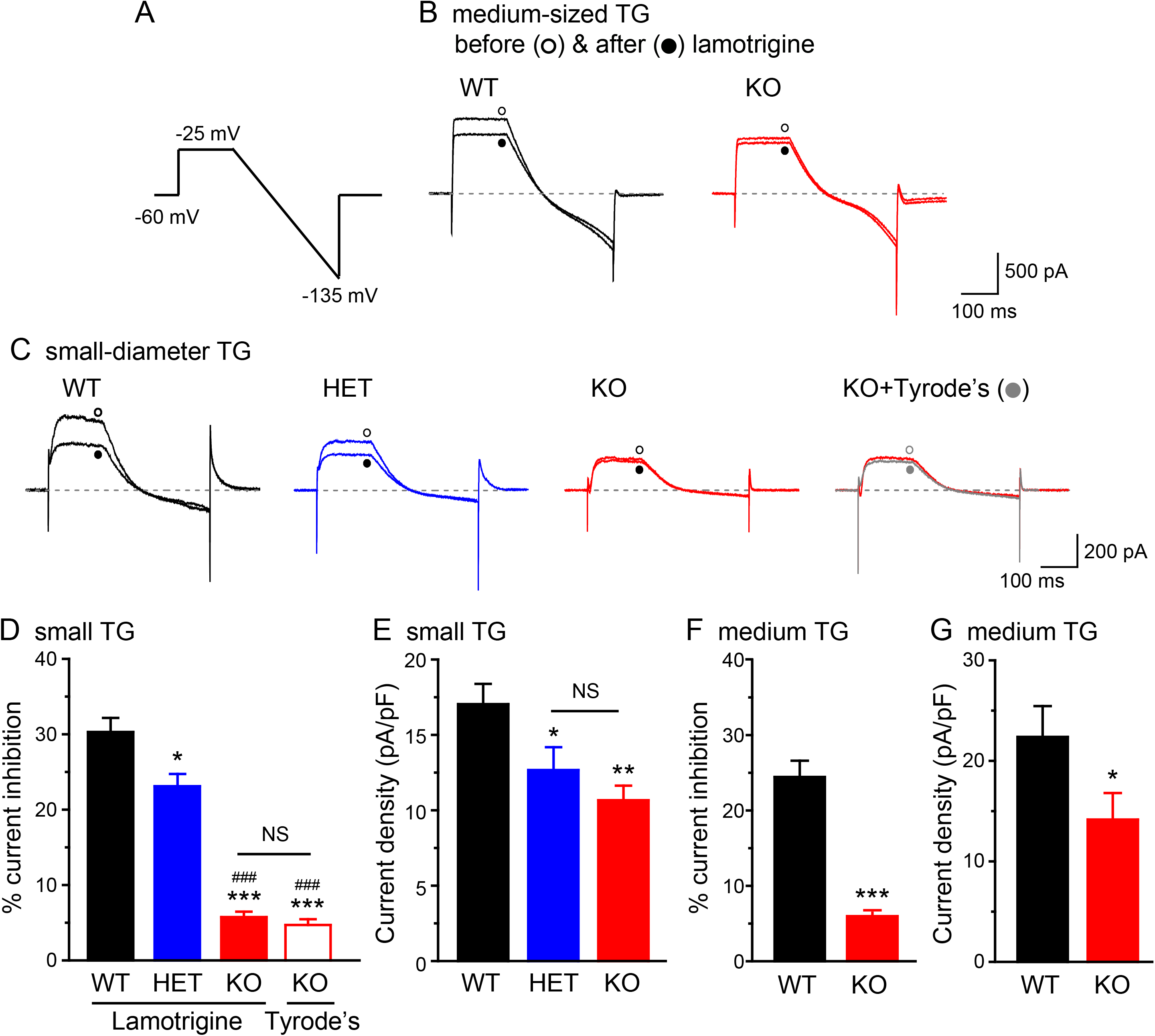
Total persistent outward currents and lamotrigine-sensitive K^+^ currents in TG neurons from WT, HET and TRESK KO mice. (A) The voltage protocol used to record persistent outward currents and to minimize transient voltage-gated K^+^, Na^+^ and Ca^2+^ currents. (B) Representative current traces from medium-sized (25–40 µm diameter) WT and KO TG neurons before and after the application of 30 µM lamotrigine. (C) Representative current traces from small-diameter (< 25 µm) WT, HET and KO TG neurons before and after the application of 30 µM lamotrigine (or Tyrode’s solution), respectively. (D-E) The percentage of lamotrigine-sensitive persistent K^+^ currents (D) and the total persistent outward current density (E) in small-diameter TG neurons (n = 14–20 neurons in each group, all measured at the end of the step to −25 mV). *p < 0.05; **p < 0.01; ***p < 0.001; one-way ANOVA with post hoc Bonferroni test, compared with the WT group; ^###^p< 0.001; compared with the HET group. NS: no statistically significant difference. (F-G) The percentage of lamotrigine-sensitive persistent K^+^ currents (F) and the total persistent K^+^ current density (G) in medium-sized TG neurons from WT and KO mice (n = 15 and 21 neurons, respectively). *p < 0.05; ***p < 0.001; two-tailed t-test.

We proceeded to investigate how loss of TRESK affects TG neuronal excitability. The mean resting membrane potential (V_rest_) was comparable between adult WT, HET and TRESK KO TG neurons (Table 1), consistent with previous studies suggesting that endogenous TRESK activity does not regulate V_rest_ in PANs (Dobler et al., 2007; Guo et al., 2014; Liu et al., 2013). To test how ubiquitous loss of TRESK current affects the excitability of TG neurons, we divided TG neurons into three subpopulations and used current-clamp recording to compare the passive and active electrophysiological properties of WT, HET and TRESK KO neurons within each group. Neurons were first sorted by soma diameter into small- (< 25 µm) and medium-sized (25– 40 µm) groups. The majority of small TG neurons are nociceptors (Harper & Lawson, 1985a, 1985b), with a small subset representing the C-low threshold mechanorecepetors (C-LTMRs) that transmit innocuous touch sensation (Lawson et al., 1997; Seal et al., 2009). Most of the medium-sized TG neurons are low-threshold mechanoreceptors with myelinated Aβ fibers (Bae et al., 2015; Goldstein et al., 1991; Perry et al., 1991). Small TG neurons were further divided into IB4-positive (IB4^+^) and IB4-negative (IB4^−^) groups, based on their ability to bind to fluorescently labeled IB4. It is well established that the small IB4^+^ and IB4^−^ PANs exhibit distinct neurochemical, anatomical, and electrophysiological properties and encode different pain modalities (Cavanaugh et al., 2009; Choi et al., 2007; Fang et al., 2006; Scherrer et al., 2009; Snider & McMahon, 1998; Stucky & Lewin, 1999). Indeed, both the rheobase value and the spike frequency were significantly different between small IB4^−^, small IB4^+^ and medium-sized TG neurons from WT mice (Figures 3A, G, H).

**Figure 3.**
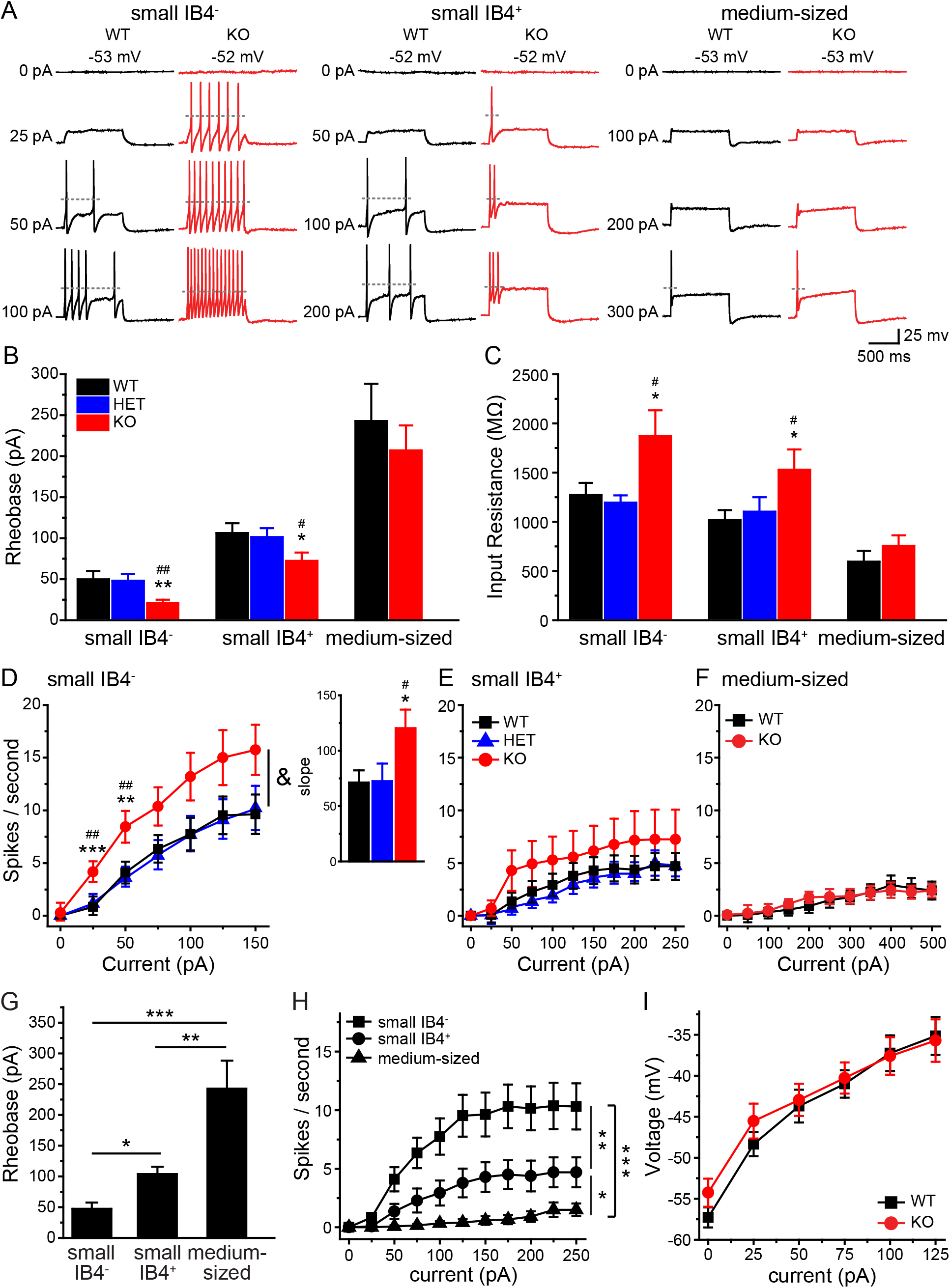
Ubiquitous loss of TRESK currents preferentially increases the excitability of small IB4^−^ TG neurons. (A) Representative traces of APs generated by incremental depolarizing current injections in TG neurons from WT and TRESK KO mice. The values of V_rest_ and the current amplitude are indicated. The dotted lines indicate membrane potential at 0 mV. (B, C) Mean rheobase (B, the minimum amount of current required to elicit at least 1 AP) and R_in_ (C) of subpopulations of TG neurons from TRESK WT, HET and KO mice (n = 14–28 neurons in each group, see Table 1 for details of the intrinsic properties of TG neurons). *p < 0.05; **p < 0.01; one-way ANOVA with post hoc Bonferroni test, compared with the corresponding WT group; ^#^p< 0.05; ^##^p < 0.01; compared with the corresponding HET group. (D-F) Input-output plots of the spike frequency in response to incremental depolarizing current injections in small IB4^−^ (D), small IB4^+^ (E) and medium-sized (F) TG neurons from TRESK WT, HET and KO mice (same neurons as in B). ^&^p < 0.05; two-way RM ANOVA; TRESK KO neurons versus WT and HET groups. **p < 0.01; ***p < 0.001; one-way ANOVA with post hoc Bonferroni test, compared with the corresponding WT group; ^##^p < 0.01; compared with the corresponding HET group. Inset: the slope (spikes/sec/nA) of the input-output relationships between 25 pA and 125 pA current injections. *p < 0.05; ^#^p < 0.05; one-way ANOVA with post hoc Bonferroni test, compared with the WT and HET groups, respectively. (G) Mean rheobase of small IB4^−^, small IB4^+^ and medium-sized TG neurons from WT mice (same neurons as in B WT groups). *p < 0.05; **p < 0.01; ***p < 0.001; one-way ANOVA with post hoc Bonferroni test. (H) Input-output plots of the spike frequency in response to incremental depolarizing current injections in subpopulations of WT TG neurons (same WT neurons as in B WT groups). *p < 0.05; **p < 0.01; ***p < 0.001; two-way RM ANOVA. (I) The plot of steady-state membrane potential versus injected current in medium-sized TG neurons from WT and KO mice (same neurons as in B medium-sized groups).

**Table 1.**
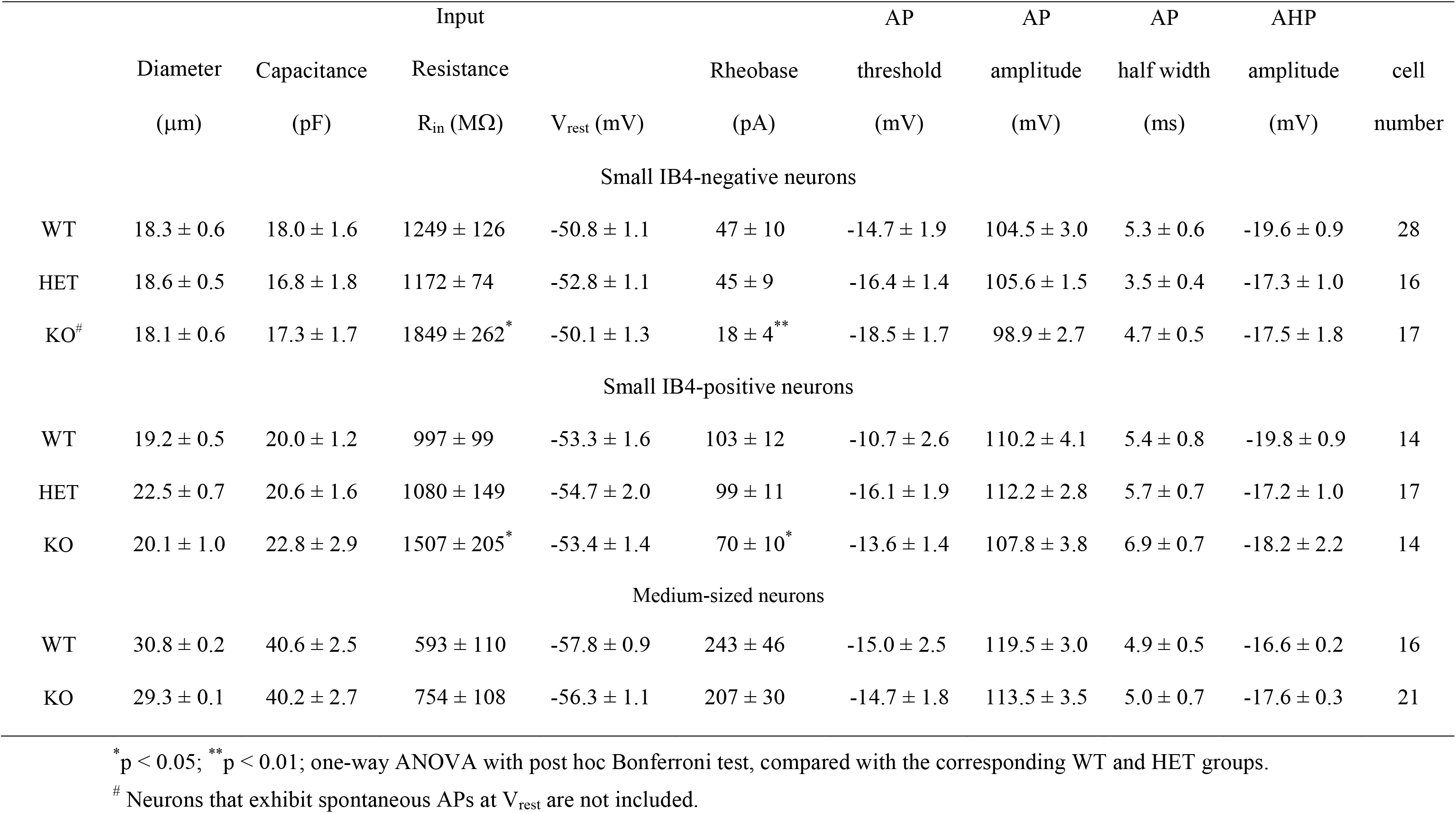
Intrinsic properties of TG neurons from adult WT, HET and TRESK KO mice.

First, we investigated whether loss of TRESK affects the excitability of small IB4^−^ TG neurons, as the majority of these neurons express CGRP, the neuropeptide that plays an important role in migraine pathophysiology (Ho et al., 2010; Price & Flores, 2007; Tao et al., 2012). Compared with WT neurons, small IB4^−^ TG neurons from KO mice exhibited a significantly higher input resistant (R_in_) and consequently a more than 50% reduction of rheobase for action potential (AP) generation (Figures 3A-C). The values of AP threshold, amplitude, half-width, or the amplitude of afterhyperpolarization (AHP) were not affected by the loss of TRESK (Table 1), suggesting that TRESK current plays a negligible role in shaping the AP waveforms.

In both WT and KO small IB4^−^ TG neurons, the number of APs initially increased almost linearly in response to incremental depolarizing current injection and plateaued upon further depolarization (Figure 3D). The slope of the input-output curve (between 25 pA and 125 pA current injections) was significantly steeper in the KO group (Figure 3D inset), and the same amount of depolarizing current evoked more APs in KO neurons than in WT (Figure 3D). We conclude that the endogenous TRESK currents control the onset and the frequency of APs in small IB4^−^ TG neurons.

Next, we compared the excitability of small IB4^+^ TG neurons from WT and KO mice. Loss of TRESK also resulted in higher R_in_ and lower rheobase in this TG subgroup (Figures 3A-C). In WT TG culture, the input-output curve of IB4^+^ neurons was much flatter than that of small IB4^−^ neurons (Figure 3H). Loss of TRESK did not significantly increase the spike frequency in small IB4^+^ TG neurons (Figure 3E), indicating that the endogenous TRESK activity regulates AP initiation but not AP frequency in this TG subpopulation.

Among the three subtypes of WT TG neurons, the medium-sized neurons had the lowest R_in_, highest rheobase and the majority of them generated a single spike in response to both threshold and supra-threshold current injections (Figures 3B-C, G-H, (Ratte et al., 2014)). Surprisingly, loss of TRESK did not alter R_in_, rheobase or spike frequency in medium-sized TG neurons at all (Figures 3A-C, F), despite the significant reduction of total persistent outward current at −25mV (Figure 2F). Of note, injection of depolarizing currents induced similar membrane potential changes between – 60 mV and −35 mV in WT and KO medium-sized TG neurons (Figure 3I), suggesting that there is little endogenous TRESK activity around V_rest_ to oppose the membrane depolarization in WT medium-sized TG neurons.

Of all the WT TG neurons we recorded (n = 214), none showed spontaneous APs at V_rest_. On the contrary, 11 TRESK KO TG neurons exhibited spontaneous APs at V_rest_, all of which belonged to the small IB4^−^ subpopulation. This accounted for 5.6% (11 of 197) of the total KO TG neurons and 9.8% (11 of 112) of KO small IB4^−^ TG neurons that we recorded for this study (Figure 4A); again indicating that loss of TRESK preferentially increases the intrinsic excitability of small IB4^−^ TG neurons. Compared with WT small IB4^−^ TG neurons, KO neurons with spontaneous APs had a similar V_rest_ (−48 ± 2 mV) but a considerably lower AP threshold (Figure 4B).

**Figure 4.**
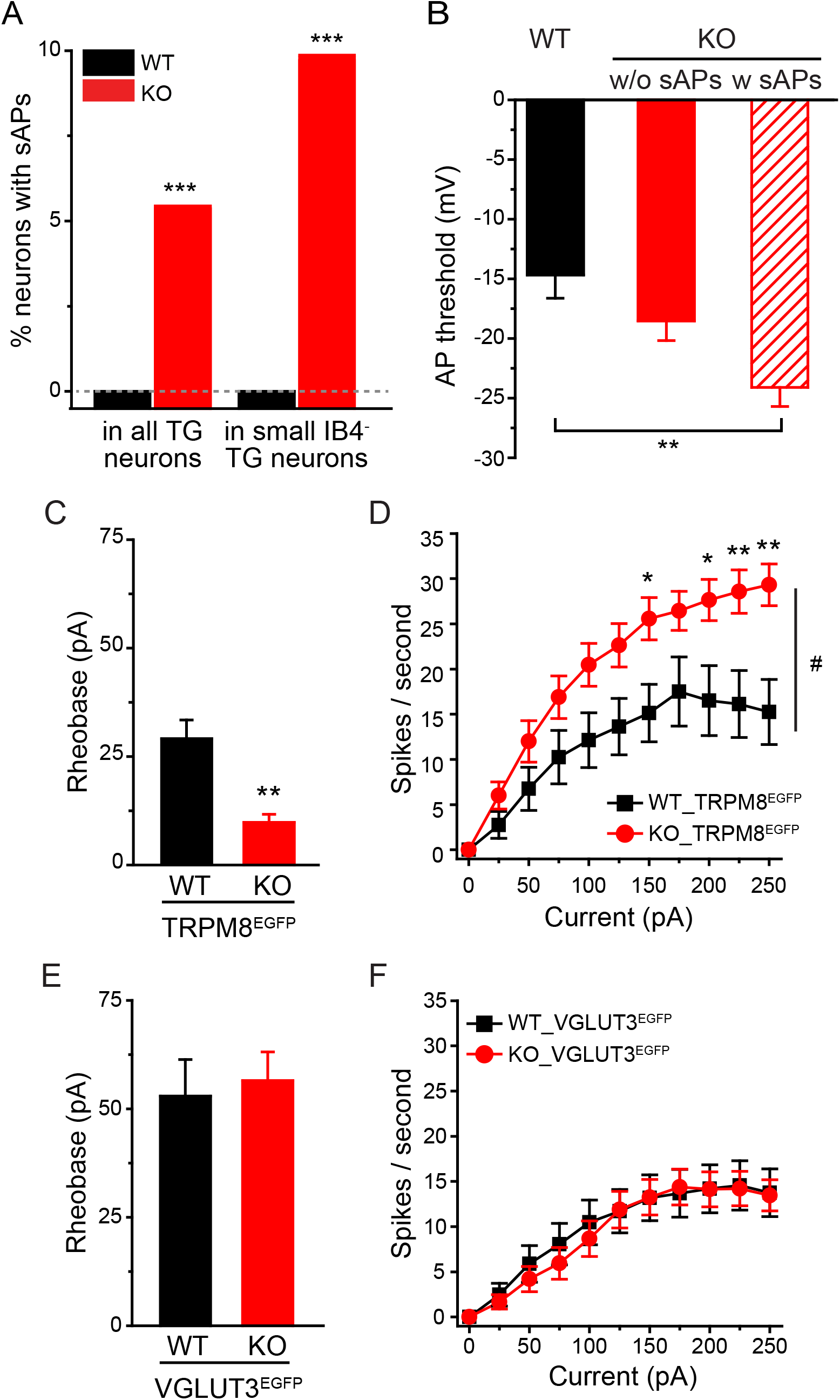
Loss of TRESK increases the excitability of small IB4^−^ TG neurons expressing TRPM8 but not those expressing VGLUT3. (A) The percentage of neurons with spontaneous APs (sAPs) in all TG neurons and in small IB4^−^ TG neurons from WT and TRESK KO mice, respectively. ***p < 0.001, Fisher’s exact test between the corresponding WT and KO groups. The dashed line indicates 0%. (B) The AP threshold of WT and KO small IB4^−^ TG neurons without spontaneous APs (w/o sAPs, same neurons as in Figure 3B) as well as KO TG neurons with spontaneous APs (w sAPs, n = 11). **p < 0.01, one-way ANOVA with post hoc Bonferroni test. (C) Mean rheobase of EGFP^+^ TG neurons from WT_TRPM8^EGFP^ and KO_TRPM8^EGFP^ mice (n = 14 and 18 neurons, respectively). **p < 0.01; two-tailed t-test. (D) The spike frequency in response to incremental depolarizing current injections in EGFP^+^ TG neurons from WT_TRPM8^EGFP^ and KO_TRPM8^EGFP^ mice (same neurons as in C). ^#^p < 0.05; two-way RM ANOVA. *p < 0.05; **p < 0.01; two-tailed t-test between the corresponding WT and KO groups. (E-F) Mean rheobase and spike frequency of EGFP^+^ TG neurons from WT_VGLUT3^EGFP^ and KO_VGLUT3^EGFP^ mice (n = 19 and 27 neurons, respectively).

We also tested whether TG neuronal excitability was altered in TRESK HET mice. Despite the reduction of both TRESK and total persistent outward currents (Figures 2B-D), neither small IB4^−^ nor IB4^+^ TG neurons from HET mice exhibited any changes of the measured passive and active electrophysiological properties (Figures 3B-E, Table 1). This is consistent with our previous finding that only a substantial reduction of the endogenous TRESK activity results in hyper-excitation of TG neurons; and a moderate decrease of TRESK current is well tolerated (Guo et al., 2014).

In addition to neurons that express CGRP, the small IB4^−^ TG subpopulation also consists of neurons that express the cold sensor TRPM8 channels and C-LTMRs that express vesicular glutamate transporter 3 (VGLUT3), respectively (Le Pichon & Chesler, 2014; Usoskin et al., 2015). These neurons were likely under-sampled when we recorded from small IB4^−^ TG neurons, as they each accounted for less than 20% of small IB4^−^ TG neurons. To identify these neurons in TG culture, we generated WT and TRESK KO mice that express enhanced green fluorescent protein (EGFP) from one TRPM8 locus (WT_TRPM8^EGFP^ and KO_TRPM8^EGFP^ mice, (Dhaka et al., 2008)) as well as mice that contain a transgenic allele that expresses EGFP under the control of the genomic sequences that regulate the expression of endogenous VGLUT3 (WT_VGLUT3^EGFP^ and KO_VGLUT3^EGFP^ mice, (Seal et al., 2009)). Compared to their WT counterparts, the EGFP-positive (EGFP^+^), TRPM8-expressing TG neurons from TRESK KO mice showed a 70% reduction of rheobase and a much higher spike frequency in response to supra-threshold current injection (Figures 4C, D). On the contrary, loss of TRESK had no effect on the excitability of EGFP^+^, VGLUT3-expressing C-LTMRs (Figures 4E, F). We also quantified the abundance of CGRP- and TRPM8-expressing TG neurons under our culture conditions. Only 11% (268 of 2365 neurons from 5 WT_TRPM8^EGFP^ mice) of small IB4^−^ TG neurons were EGFP^+^; whereas 73% (459 of 626 neurons from 3 WT mice) of them contained CGRP-immunoreactivity. This indicates that the results obtained from small IB4^−^ TG neurons largely reflected changes consistent with how loss of TRESK affects the excitability of CGRP-expressing (CGRP^+^) PANs.

Having characterized how loss of TRESK affects TG neuronal excitability, we proceeded to investigate whether loss of TRESK affects the excitability of small-diameter DRG neurons. As in TG neurons, 30 µM lamotrigine blocked 20-30% of persistent outward currents in every adult WT DRG neurons that we recorded but had no significant effect on currents in KO DRG neurons (Figure 5A). Surprisingly, the size of total persistent outward current remained comparable between WT and KO DRG neurons (Figure 5B), suggesting that loss of endogenous TRESK current is fully compensated in DRG neurons. Consequently, in either small IB4^−^ or small IB4^+^ DRG subpopulation, WT and KO neurons exhibit similar R_in_, rheobase, spike frequency as well as other passive and active electrophysiological properties (Figures 5C-F, Table 2). Our findings are inconsistent with a previous study reporting a reduction of total persistent outward currents and AP rheobase in DRG neurons from mice expressing non-functional TRESK channels (Dobler et al., 2007). More work is needed to resolve the discrepancy. It is possible that results differ based on the duration of time neurons are maintained *in vitro* after dissociation. Here, we recorded DRG neurons between 2-4 days *in vitro* (DIV), whereas in earlier work neurons were recorded between 11-14 DIV. It has been reported that WT DRG neurons cultured for 5 DIV exhibited altered K^+^ channel expression and excitability (Dawes et al., 2018; Martinez-Espinosa et al., 2015).

**Figure 5.**
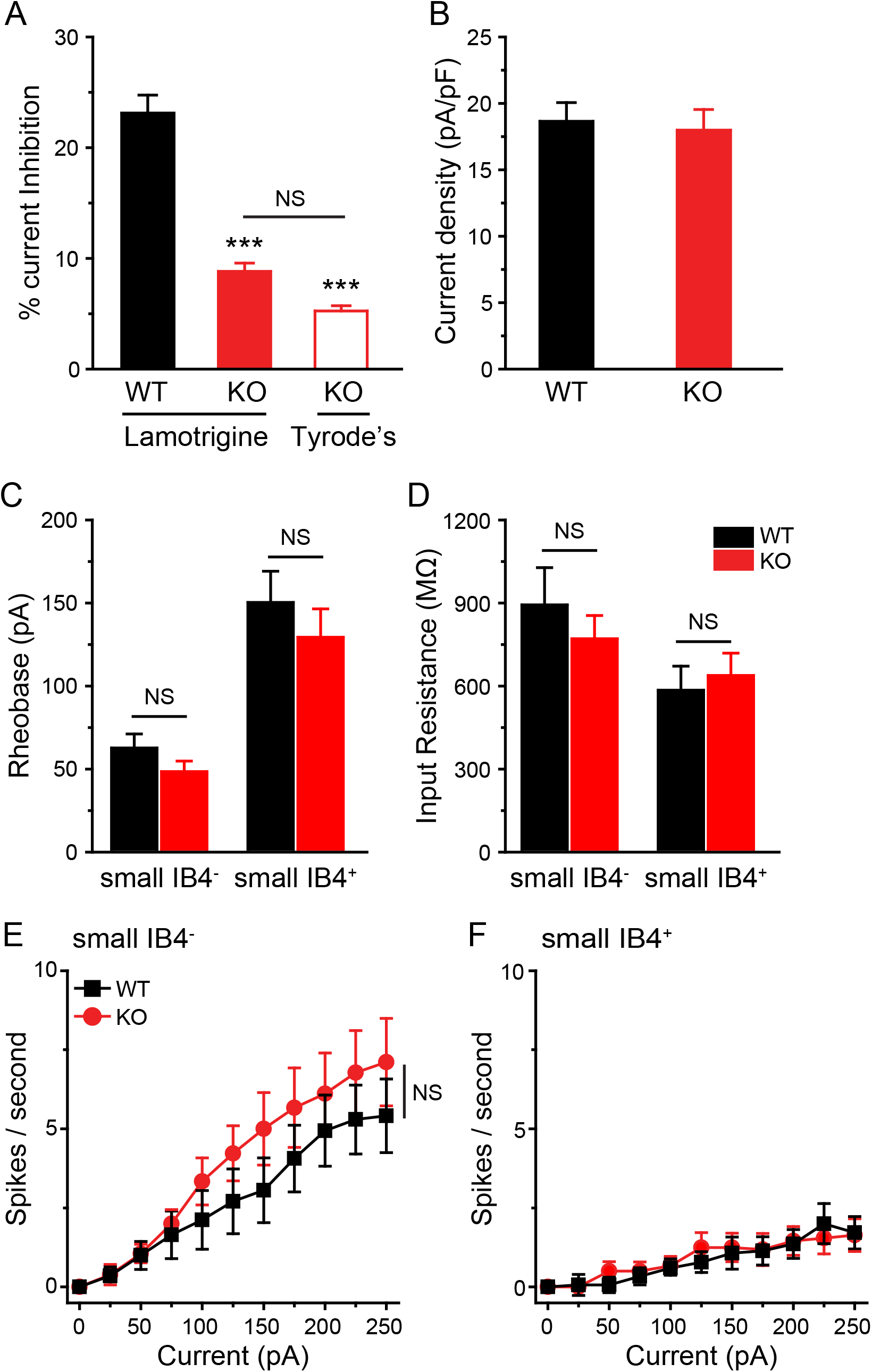
Loss of TRESK does not alter the excitability of small lumbar DRG neurons. (A-B) The percentage of lamotrigine-sensitive persistent K^+^ currents (A) and the total persistent outward current density (B) in small-diameter DRG neurons from adult WT and TRESK KO mice (n = 14–20 neurons in each group). ***p < 0.001; one-way ANOVA with post hoc Bonferroni test, compared with the WT group. NS: no statistically significant difference. (C-D) Mean rheobase (C) and R_in_ (D) of small IB4^−^ and IB4^+^ DRG neurons from WT and TRESK KO mice (n = 16–19 neurons in each group, see Table 2 for details of the intrinsic properties of DRG neurons). (E-F) Input-output plots of the spike frequency in response to incremental depolarizing current injections in small IB4^−^ (E) and small IB4^+^ (F) DRG neurons from WT and KO mice (same neurons as in C).

**Table 2.**
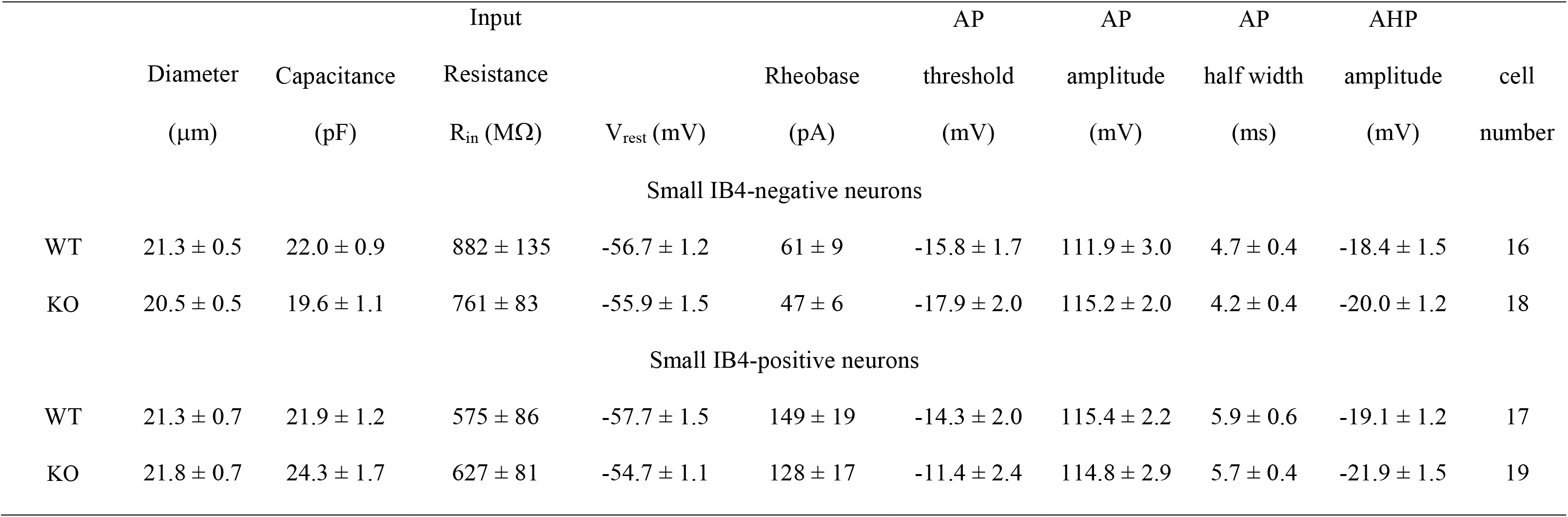
Intrinsic properties of small-diameter DRG neurons from adult WT and TRESK KO mice.

Taken together, we conclude that ubiquitous loss of TRESK in all TG and DRG neurons preferentially increases the intrinsic excitability of small IB4^−^ TG nociceptors that express CGRP and/or TRPM8, suggesting that the contribution of endogenous TRESK channel to PAN excitability is cell-type specific and is not determined by its expression pattern.

### Loss of TRESK selectively increases the number of capsaicin-responsive (Cap^+^) neurons in the IB4^−^ TG subpopulation

Here, we investigated whether loss of TRESK affect the responses of TG neurons to noxious stimuli, for example, exposure to capsaicin that selectively activates TRPV1 channels. We applied capsaicin (100 nM) to small-diameter TG neurons in culture and recorded their responses under current-clamp. In TG culture from adult WT mice, capsaicin elicited depolarization in 9 of 25 (36%) of neurons, 6 of which also exhibited multiple spikes (Figures 6A, B black). The percentage of small TG neurons that responded to capsaicin was significantly higher in culture from KO mice (67%, 24 of 36). In 18 of 36 KO TG neurons, capsaicin evoked both depolarization and multiple spikes. An additional 6 (17%) KO neurons exhibited capsaicin-induced depolarization only (Figure 6B red). The higher percentage of KO neurons with capsaicin-evoked spikes likely results from the increase in intrinsic excitability. However, that capsaicin depolarized more KO neurons than WT suggests that loss of TRESK may also increase the number of TG neurons exhibiting functional TRPV1 channels.

**Figure 6.**
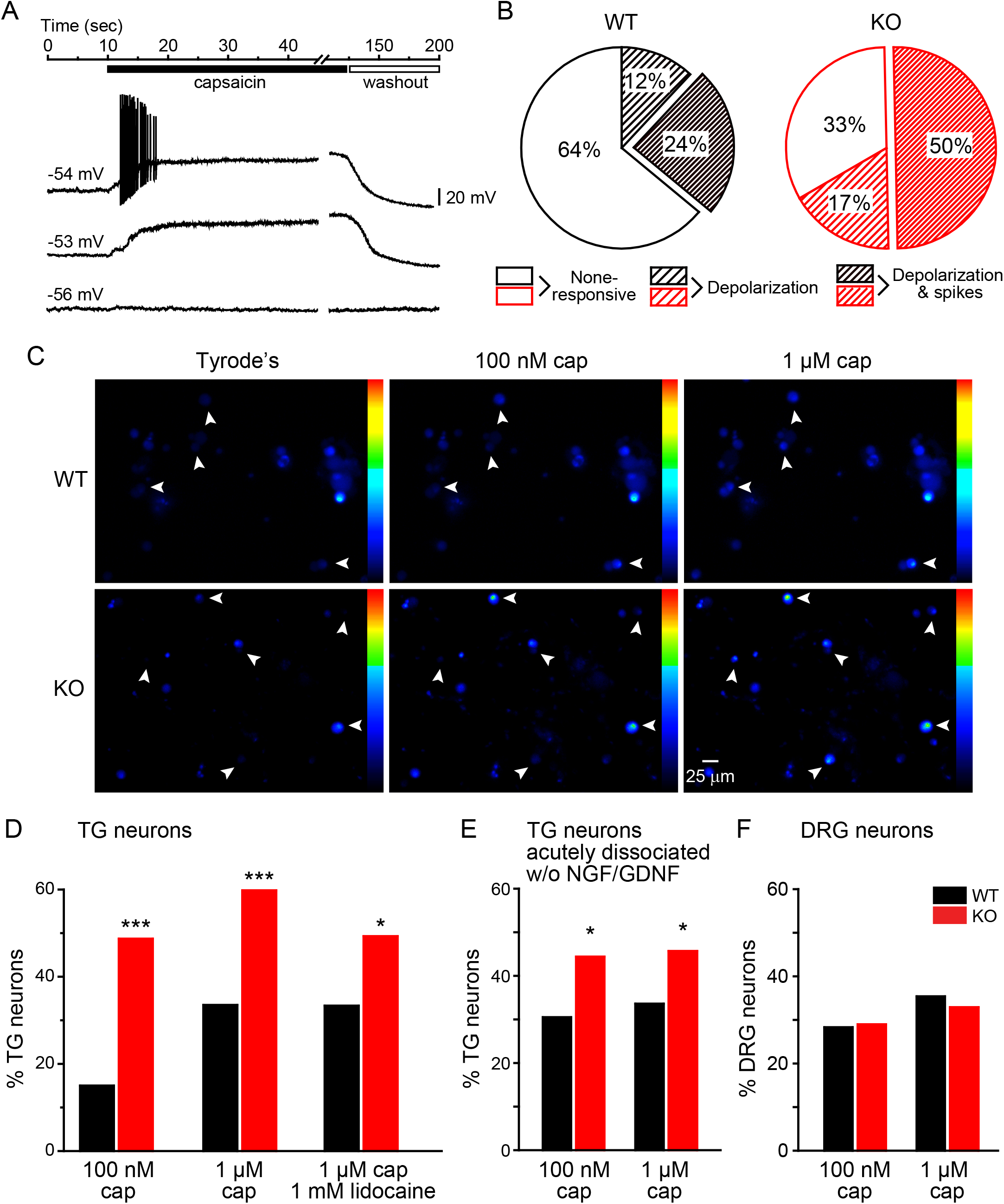
The number of Cap^+^ neurons is increased in cultured TG but not DRG neurons from TRESK KO mice. (A) Representative current-clamp traces of 3 small-diameter WT TG neurons in response to 100 nM capsaicin. The values of V_rest_ are indicated. (B) The percentage of WT and KO TG neurons that exhibit depolarization, burst of spikes or no response to 100 nM capsaicin (n = 25 WT and 36 KO neurons). p < 0.05; χ^2^ test between WT and KO groups. (C) Representative Fura-2 images of cultured TG neurons from WT and KO mice before and after applications of 100 nM and 1 µM capsaicin, respectively. The blue to green color transition reflects increase in intracellular Ca^2+^. The arrowheads indicate neurons that respond to capsaicin. (D) The percentage of WT and KO TG neurons that responds to 100 nM and 1 µM capsaicin (cap, n = 146 WT and 117 KO TG neurons from 3 experiments), respectively. In another set of experiment, the extracellular solutions contained 1 mM lidocaine (n = 111 WT and 138 KO neurons). *p < 0.05; ***p < 0.001; Fisher’s exact test between the corresponding WT and KO groups. (E) The percentage of Cap^+^ neurons in acutely dissociated WT and KO TG neurons without NGF and GDNF exposure (n = 131 WT and 153 KO neurons from 3 experiments). (F) The percentage of WT and KO DRG neurons that responds to 100 nM and 1 µM capsaicin (n = 628 WT and 490 KO DRG neurons from 3 experiments), respectively.

To test this possibility, we used the ratiometric Ca^2+^ indicator fura-2 to measure capsaicin-induced Ca^2+^ influx in TG neurons from WT and TRESK KO mice. This allows us to compare the responses to capsaicin in large numbers of neurons (Figure 6C). Consistent with the electrophysiology data, the percentage of KO neurons that responded to capsaicin (0.1 to 1 µM, > 20% increase from baseline) was significantly higher than that of the WT group (Figures 6C-D). To test whether this results from prolonged exposure of TG neurons to nerve growth factor (NGF) and/or glial cell line-derived neurotrophic factor (GDNF) *in vitro*, we repeated the experiment in WT and KO TG neurons maintained in media without NGF and GDNF within 24 hours after dissociation. Once again, the fraction of Cap^+^ KO TG neurons was significantly higher than that of the WT group (Figure 6E). In contrast, the percentage of Cap^+^ neurons was similar in DRG cultures from WT and KO mice (Figure 6F), indicating that genetic loss of TRESK selectively increases the number of Cap^+^ neurons in TG.

In another control experiment, we measured capsaicin-induced Ca^2+^ influx with 1 mM lidocaine in the extracellular solution. This served two purposes. First, this allowed us to measure Ca^2+^ influx exclusively through TRPV1 channels, as lidocaine blocks all voltage-gated Na^+^ channels, thereby eliminating capsaicin-evoked APs and the subsequent AP-induced Ca^2+^ influx through voltage-gated Ca^2+^ channels. Second, lidocaine at this concentration blocks more than 70% of currents through mouse TRESK channels without affecting TRPV1 currents (Keshavaprasad et al., 2005; Leffler et al., 2008; Sano et al., 2003), allowing us to test whether transient reduction of TRESK activity affects capsaicin-induced Ca^2+^ influx in TG neurons. After eliminating AP-induced Ca^2+^ influx, the percentage of Cap^+^ neurons was still higher in TG culture from TRESK KO mice (Figure 6D). Of note, the fraction of Cap^+^ neurons in WT TG culture was not altered by lidocaine (Figure 6D), suggesting that genetic loss of TRESK, but not transient reduction of TRESK activity, increases the number of TG neurons exhibiting functional TRPV1 channels.

We also compared basal and capsaicin-evoked Ca^2+^ influx in individual TG neurons. Basal Ca^2+^ level was similar between WT and KO neurons (Figure 7A), indicating that loss of TRESK does not affect resting Ca^2+^ concentration. The magnitude of peak capsaicin-evoked Ca^2+^ influx was also comparable between WT and KO TG neurons (Figure 7B). Taken together, these results suggest that genetic loss of TRESK results in the expression of functional TRPV1 channels in a subpopulation of TG neurons that normally do not respond to capsaicin. However, loss of TRESK does not further enhance TRPV1 activity in TG neurons that normally respond to capsaicin.

**Figure 7.**
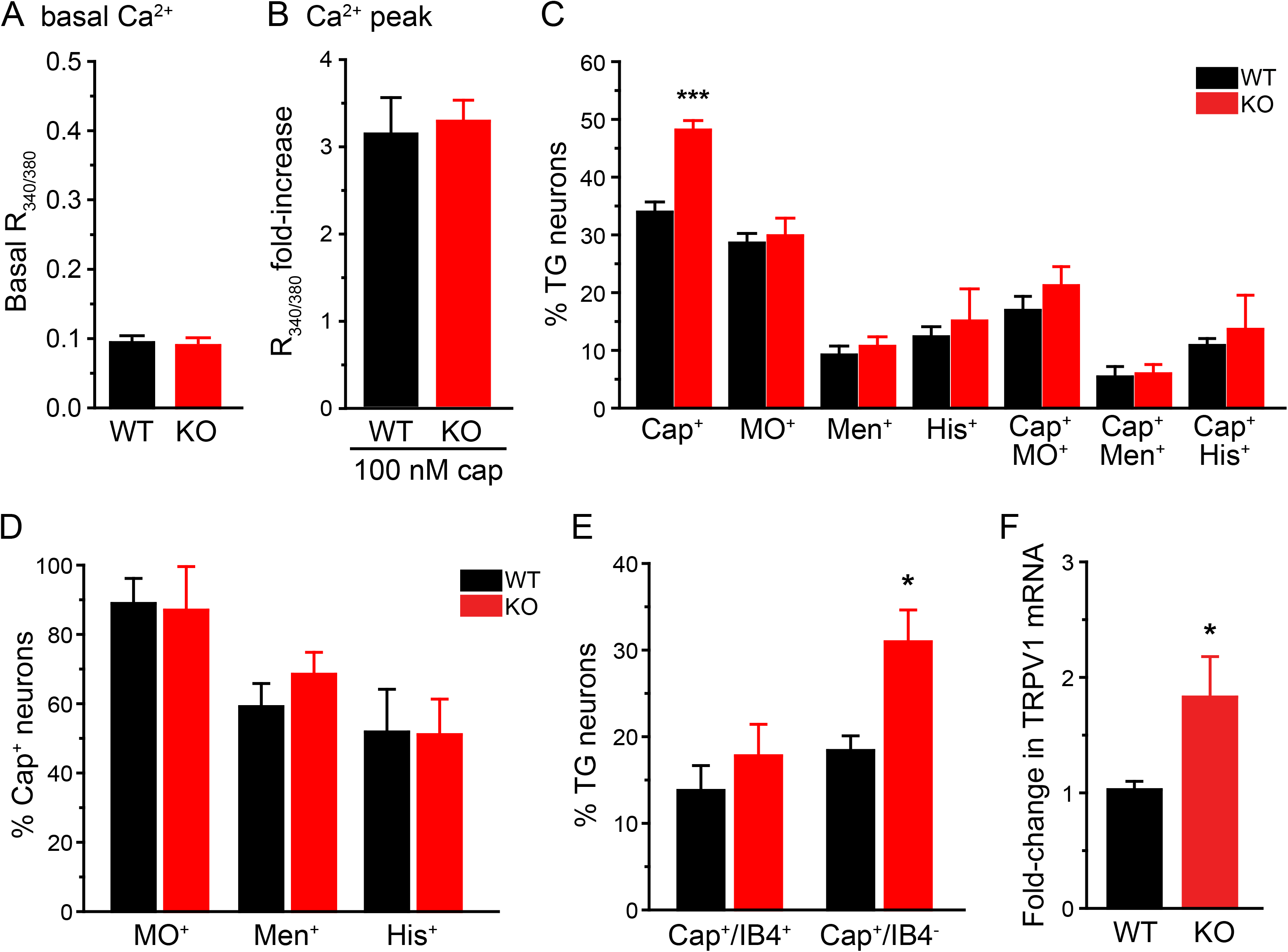
The percentage of Cap^+^ neurons is selectively increased in the small IB4^−^ TG subpopulation from TRESK KO mice. (A) Basal R_340/380_ values of WT and KO TG neurons (n = 146 WT and 117 KO neurons from 3 experiments). (B) Average fold change of peak R_340/380_ values ([peak_R_340/380_ – basal_R_340/380_] / basal_R_340/380_) in WT and KO TG neurons that respond to 100 nM capsaicin (n = 58 WT and 72 KO neurons from 3 experiments). (C) The percentage of WT and KO TG neurons that responds to 1 µM capsaicin (Cap^+^), 100 µM mustard oil (MO^+^), 100 µM menthol (Men^+^), 100 µM histamine (His^+^) as well as neurons that are Cap^+^MO^+^, Cap^+^Men^+^ and Cap^+^His^+^, respectively (n = 1010 WT and 940 KO TG neurons from 10 experiments). ***p < 0.001; two-tailed t-test between the corresponding WT and KO groups. (D) The percentage of Cap^+^ neurons in MO^+^, Men^+^ and His^+^ TG subpopulations in WT and KO culture, respectively (same neurons as in C). (E) The percentage of WT and KO TG neurons that are Cap^+^IB4^+^ and Cap^+^IB4^−^, respectively (same neurons as in C). *p < 0.05, two-tailed t-test between the corresponding WT and KO groups. (F) Relative TRPV1 mRNA expression levels in WT and KO TG (n = 10 WT and 9 KO mice). The abundance of TRPV1 mRNA was normalized to that of β-actin in individual samples. *p < 0.05, two-tailed t-test.

We next investigated whether TRESK KO TG neurons are more responsive to other chemical stimuli than WT neurons. The percentage of neurons that exhibited Ca^2+^ influx in response to 100 µM mustard oil (MO^+^), menthol (Men^+^) and histamine (His^+^) were comparable between WT and KO TG culture (Figure 7C), indicating that loss of TRESK preferentially enlarges the Cap^+^ TG population. We also sequentially measured Ca^2+^ influx evoked by capsaicin and other chemical stimuli in individual TG neurons. The percentage of TG neurons that were Cap^+^MO^+^, Cap^+^Men^+^ and Cap^+^His^+^ were similar in WT and KO TG cultures (Figure 7C). Approximately 90% of MO^+^, 60% of Men^+^ and 50% of His^+^ TG neurons were Cap^+^ in both WT and KO TG culture (Figure 7D), suggesting that loss of TRESK increases functional TRPV1 channels in TG neurons that are not activated by mustard oil, menthol or histamine.

Does the increase in functional TRPV1 occur in IB4^+^ or IB4^−^ KO TG neurons? The percentage of IB4^+^ neurons was comparable between WT and KO TG cultures (Figure 1D). Approximately 15% of TG neurons were Cap^+^IB4^+^ in both WT and KO culture (Figure 7E). On the contrary, the percentage of Cap^+^IB4^−^ TG neurons was significantly higher in TRESK KO culture than in WT culture (Figure 7E), indicating that genetic loss of TRESK preferentially increases the number of IB4^−^ TG neurons that express functional TRPV1 channels.

Lastly, we compared the level of TRPV1 mRNA in WT and KO TG tissues by quantitative polymerase chain reaction (qPCR). The relative level of TRPV1 mRNA was significantly higher in TRESK KO TG than in WT, an almost 2-fold increase (Figure 7F), indicating that genetic loss of TRESK results in upregulation of TRPV1 gene transcription in TG neurons.

### TRPV1 protein expression is increased in subpopulations of TG neurons in TRESK KO mice

We stained TG sections from WT and TRPV1 KO mice with the TRPV1 antibody. TRPV1-immunoreactivity (TRPV1-ir) was present in many small-diameter neurons in WT TG but was completely absent in TG sections from TRPV1 KO mice (Figure 8A), validating the specificity of the antibody. On average, the abundance of TRPV1-expressing (TRPV1^+^) neurons was 30% higher in TRESK KO TGs than in WT controls (Figures 8B, C), indicating that the increase in Cap^+^ neurons KO TG results from a larger population of TG neurons expressing TRPV1 proteins. The size distributions of TRPV1^+^ TG neurons in WT and KO mice were similar (Figure 8E, F), the majority (> 80%) of them had a cross-sectional area smaller than 500 μm^2^ and belonged to small-diameter (< 25 µm) PANs.

**Figure 8.**
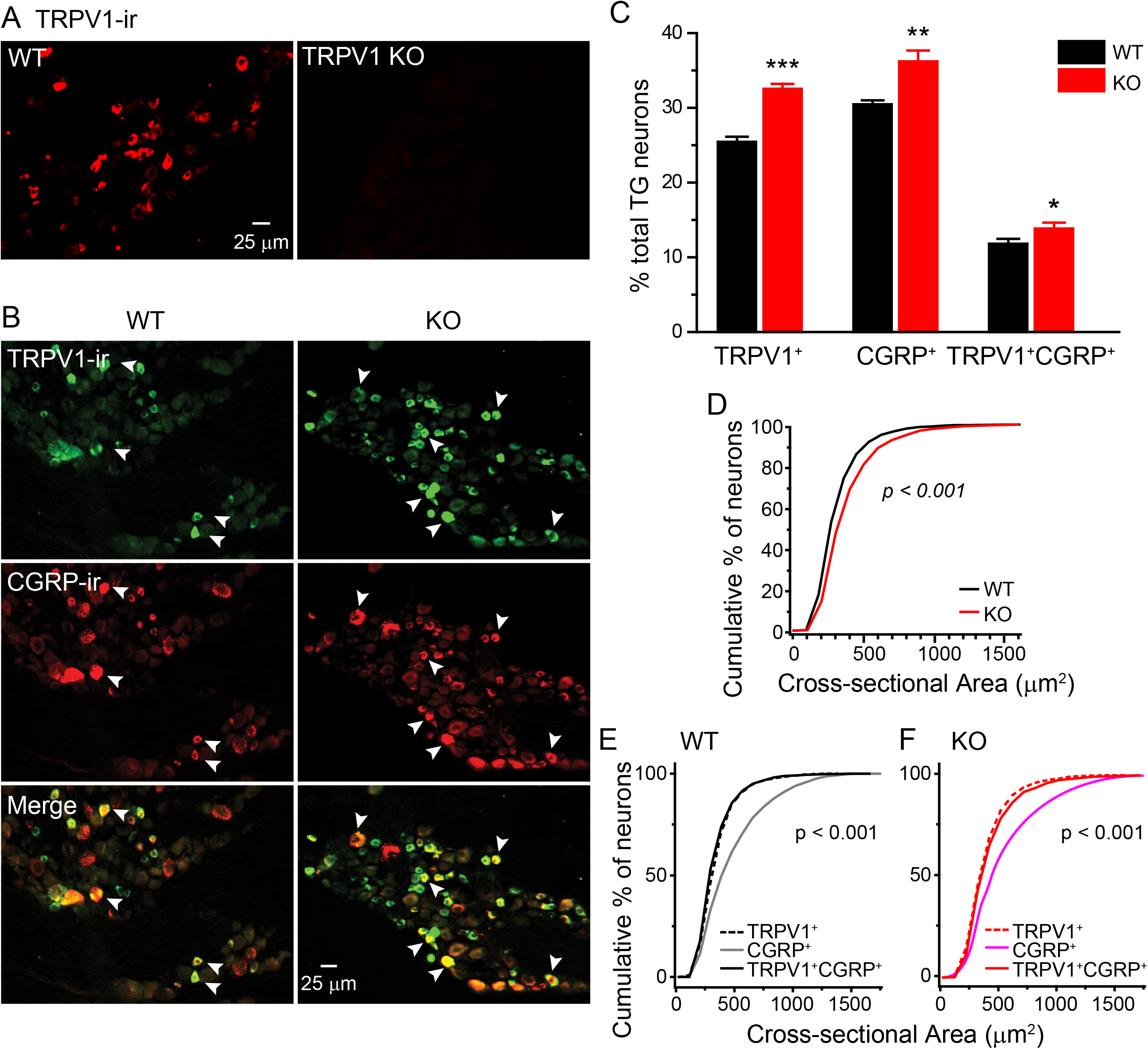
Loss of TRESK increases the number of TG neurons expressing CGRP and TRPV1. (A) Representative images of TRPV1-ir in TG sections from WT and TRPV1 KO mice. TRPV1-ir is present in some small TG neurons in the WT section. There is no staining in the TRPV1 KO TG section, validating the specificity of the TRPV1 antibody. (B) Representative images of TG sections from WT and TRESK KO mice double stained with antibodies against TRPV1 and CGRP. Arrowheads indicate neurons that are both TRPV1^+^ and CGRP^+^. (C) The percentage of TRPV1^+^, CGRP^+^, and TRPV1^+^CGRP^+^ TG neurons in WT and KO mice (n = 3 mice in each group; on average, 7629 TG neurons/mouse were counted). *p < 0.05; **p < 0.01; ***p < 0.001; two-tailed t-test between the corresponding WT and KO groups. (D) Cumulative distributions of the cross-sectional areas of WT and KO TRPV1^+^CGRP^+^ TG neurons. p < 0.001; Mann–Whitney U test. (E) Cumulative distributions of the cross-sectional areas of TRPV1^+^, CGRP^+^, and TRPV1^+^CGRP^+^ neurons in WT TG (n = 3395, 3760, and 1808 neurons pooled from 3 mice, respectively, same as in C). p < 0.001 between CGRP^+^ neurons and the other groups, Kruskal-Wallis ANOVA with Dunn’s post hoc test. (F) Cumulative distributions of the cross-sectional areas of TRPV1^+^, CGRP^+^, and TRPV1^+^CGRP^+^ neurons in KO TG (n = 3175, 2685, and 1498 neurons pooled from 3 mice, respectively, same as in C). p < 0.001 between CGRP^+^ neurons and the other groups.

We went on to identify the TG subpopulation(s) in which TRPV1 expression was upregulated in KO mice. Since the increase in Cap^+^ neurons mainly occurs in the IB4^−^ KO TG neurons (Figure 7E), we co-stained TG sections for TRPV1-ir and markers that subdivide the IB4^−^ PANs (Usoskin et al., 2015). First, we examined TRPV1-ir in TG neurons expressing neuropeptide CGRP. The abundance of CGRP^+^ and TRPV1^+^CGRP^+^ neurons were both significantly increased in TRESK KO TG compared to WT (Figures 8B, C). The soma sizes of TRPV1^+^CGRP^+^ neurons in KO TG were slightly larger than those in WT (Figure 8D), but the majority (> 80%) of them still had a cross-sectional area smaller than 500 μm^2^ (Figures 8D-F).

Secondly, we quantified the overlap between TRPV1-ir and neurofilament 200-immunoreactivity (NF200-ir), a marker for myelinated PANs. In both WT and KO TG, about 35% neurons contained NF200-ir (NF200^+^), with similar soma size distributions (Figures 9A, B, E, F). Only 3.4 ± 0.3% of TG neurons were TRPV1^+^NF200^+^ in WT mice, whereas the number of TRPV1^+^NF200^+^ neurons was doubled in KO TG (7.2 ± 0.2%, Figure 9B). This accounted for about 50% of the expansion of TRPV1^+^ population in KO TG. The percentage of TRPV1^+^ neurons in NF200^+^ population and the percentage of NF200^+^ neurons in the TRPV1^+^ population were both significantly increased in KO TG (Figure 9D). The number of TRPV1^+^ neurons was increased in both small- and medium-sized (cross sectional area > 500 µm^2^) NF200^+^ TG populations in KO mice (Figures 9E, F). Overall, the soma sizes of TRPV1^+^NF200^+^ neurons in KO TG were slightly smaller than those in WT (Figure 9C).

**Figure 9.**
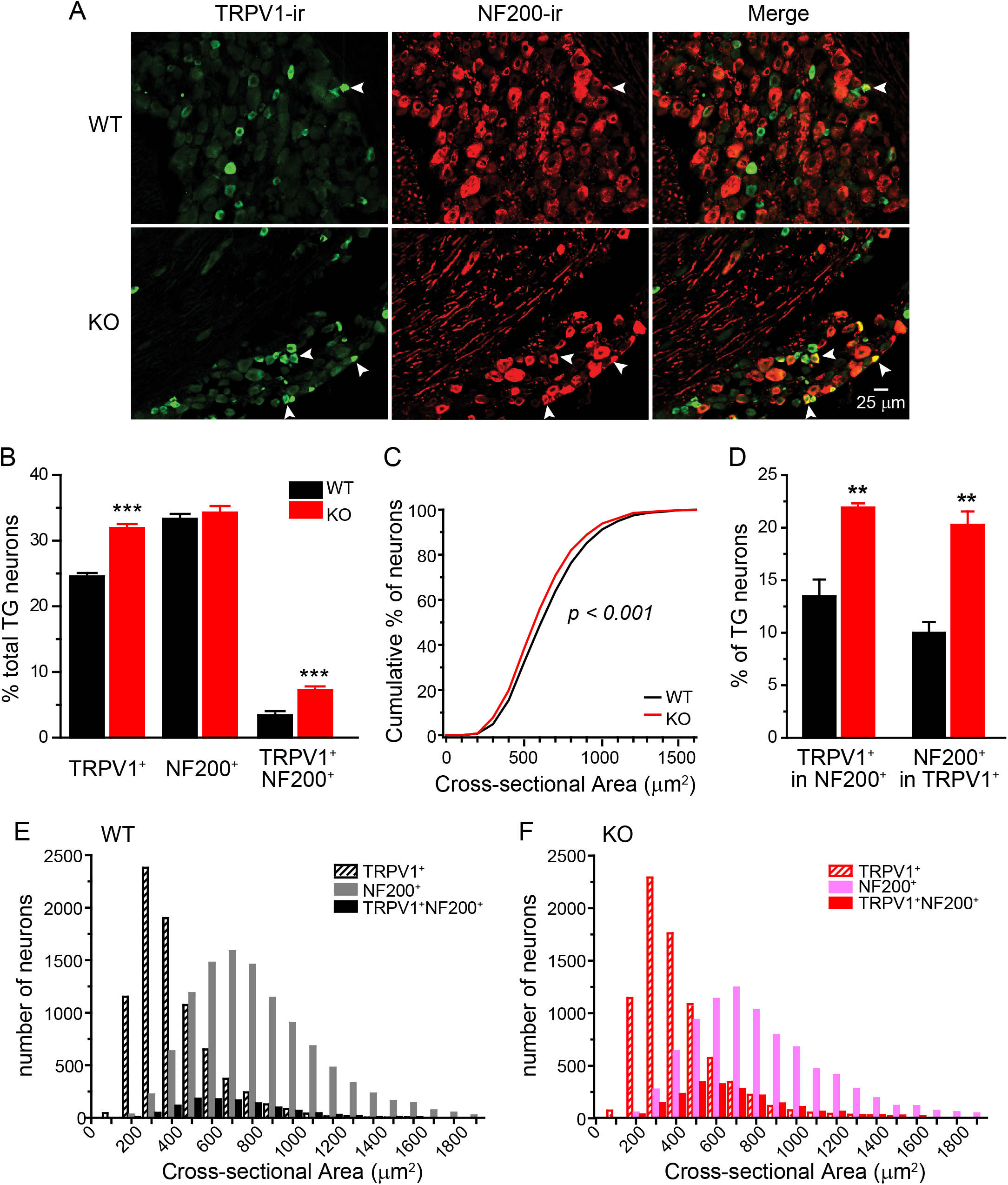
Overlap between TRPV1^+^ and NF200^+^ neurons in WT and KO TG. (A) Representative images of WT and TRESK KO TG sections double stained with antibodies against TRPV1 and NF200. Arrowheads indicate neurons that are both TRPV1^+^ and NF200^+^. (B) The percentage of TRPV1^+^, NF200^+^, and TRPV1^+^NF200^+^ TG neurons in WT and KO mice (n = 3 mice in each group; on average, 9630 TG neurons/mouse were counted). ***p < 0.001, two-tailed t-test between the corresponding WT and KO groups. (C) Cumulative distributions of the cross-sectional areas of WT and KO TRPV1^+^NF200^+^ TG neurons. p < 0.001; Mann–Whitney U test. (D) The percentage of TRPV1^+^ neurons in NF200^+^ TG population and the percentage of NF200^+^ neurons in TRPV1^+^ TG population in WT and KO mice (same mice as in B). **p < 0.01; two-tailed t-test between the corresponding WT and KO groups. (E) Histogram of soma size distributions of TRPV1^+^, NF200^+^, and TRPV1^+^NF200^+^ neurons in WT TG (n = 8081, 10789, and 1068 neurons pooled from 3 mice, respectively, same mice as in B). p < 0.001 between individual groups, Kruskal-Wallis ANOVA with Dunn’s post hoc test. (F) Histogram of soma size distributions of TRPV1^+^, NF200^+^, and TRPV1^+^NF200^+^ neurons in KO TG (n = 7663, 8174, and 1723 neurons pooled from 3 mice, respectively, same mice as B). p < 0.001 between individual groups, Kruskal-Wallis ANOVA with Dunn’s post hoc test.

We also compared TRPV1 expression in unmyelinated C-LTMRs in WT_VGLUT3^EGFP^ and KO_VGLUT3^EGFP^ mice. The percentage of EGFP^+^, VGLUT3-expressing neurons was comparable between WT and KO TG (Figure 10B). There was little overlap between the TRPV1^+^ and EGFP^+^ populations in either WT or KO TG (Figures 10A, B), indicating that loss of TRESK does not increase TRPV1 expression in TG C-LTMRs.

**Figure 10.**
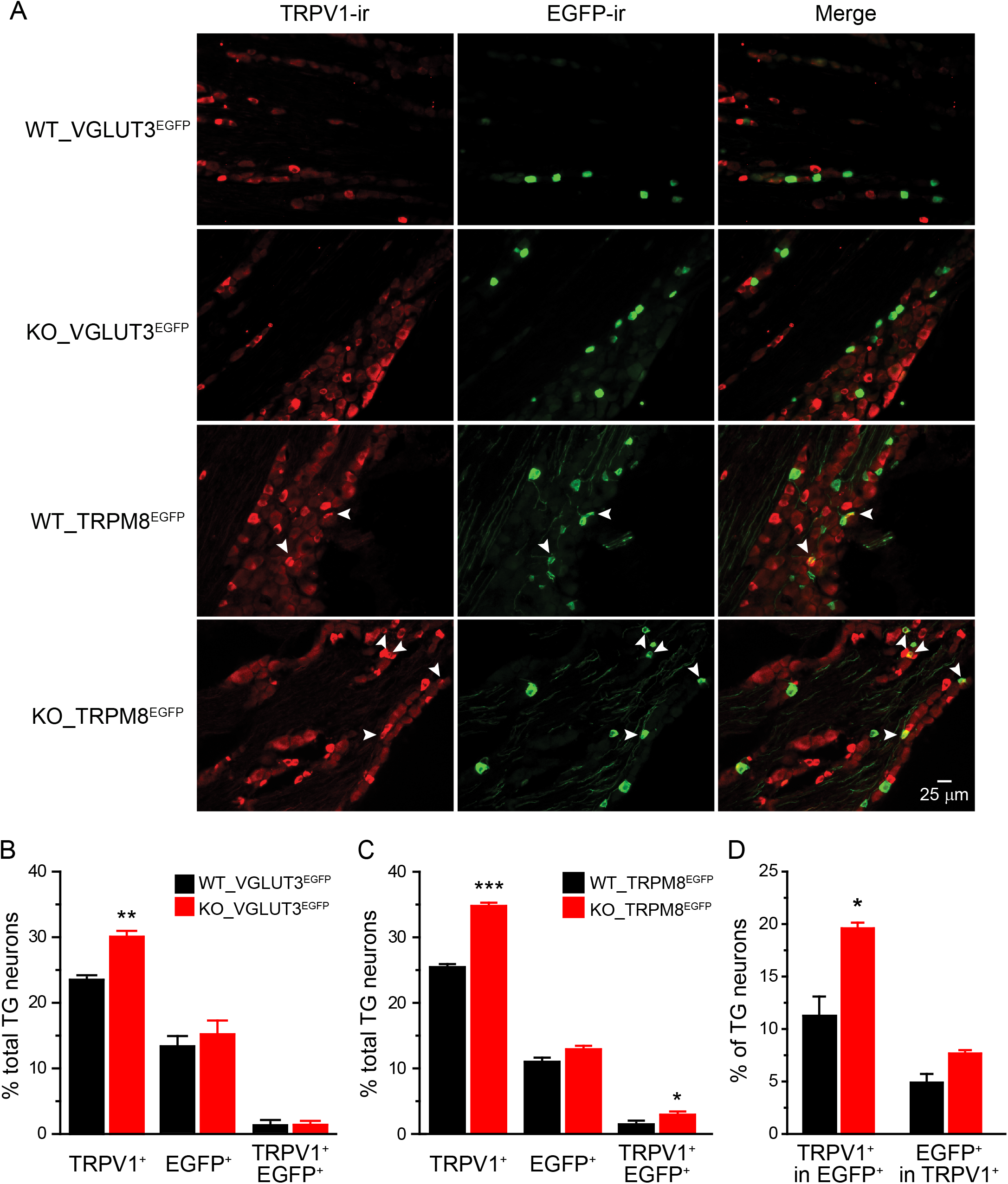
TRPV1^+^ neurons are increased in KO TG neurons expressing TRPM8 but not those expressing VGLUT3. (A) Representative images of TG sections from WT_VGLUT3^EGFP^, KO_VGLUT3^EGFP^, WT_TRPM8^EGFP^ and KO_TRPM8^EGFP^ mice double stained with antibodies against TRPV1 and EGFP. Arrowheads indicate neurons that are both TRPV1^+^ and EGFP^+^. (B) The percentage of TRPV1^+^, EGFP^+^, and TRPV1^+^EGFP^+^ TG neurons in WT_VGLUT3^EGFP^ and KO_VGLUT3^EGFP^ mice (n = 3 mice in each group; on average, 6845 TG neurons/mouse were counted). **p < 0.01; two-tailed t-test. (C) The percentage of TRPV1^+^, EGFP^+^, and TRPV1^+^EGFP^+^ TG neurons in WT_TRPM8^EGFP^ and KO_TRPM8^EGFP^ mice (n = 4 mice in each group; on average, 12027 TG neurons/mouse were counted). *p < 0.05; ***p < 0.001; two-tailed t-test. (D) The percentage of TRPV1^+^ neurons in EGFP^+^ TG population and the percentage of EGFP^+^ neurons in TRPV1^+^ TG population in WT_TRPM8^EGFP^ and KO_TRPM8^EGFP^ (same mice as in C). *p < 0.05; two-tailed t-test.

Lastly, we investigated whether loss of TRESK leads to expansion of TRPV1 expression in TG neurons expressing TRPM8 channels. In sections from WT_TRPM8^EGFP^ and KO_TRPM8^EGFP^ mice, we found that the percentage of EGFP^+^, TRPM8-expressing neurons was similar between WT and KO TGs (Figure 10C). The abundance of TPPV1^+^EGFP^+^ populations was significantly increased in KO TG compared to WT (Figure 10C). The percentage of TRPV1^+^ neurons in EGFP^+^ population was doubled in KO TG (Figure 10D). Taken together, in the absence of TRESK, TRPV1 channel is expressed in more IB4^−^ TG neurons that express CGRP, NF200 and/or TRPM8 but not VGLUT3, suggesting that genetic loss of TRESK may convert some mechano- or cold-sensitive TG neurons to polymodal nociceptors.

### Loss of TRESK preferentially enhances trigeminal nociception

TRESK KO mice were grossly normal. Both male and female KO mice gained weight normally. The performance of WT and KO mice did not differ in open field and rotarod tests (Figures 11A-G), indicating that loss of TRESK does not affect general locomotion, coordination or the level of anxiety in mice. This allowed us to compare the nociceptive responses of WT and TRESK KO mice in a battery of pain models.

**Figure 11.**
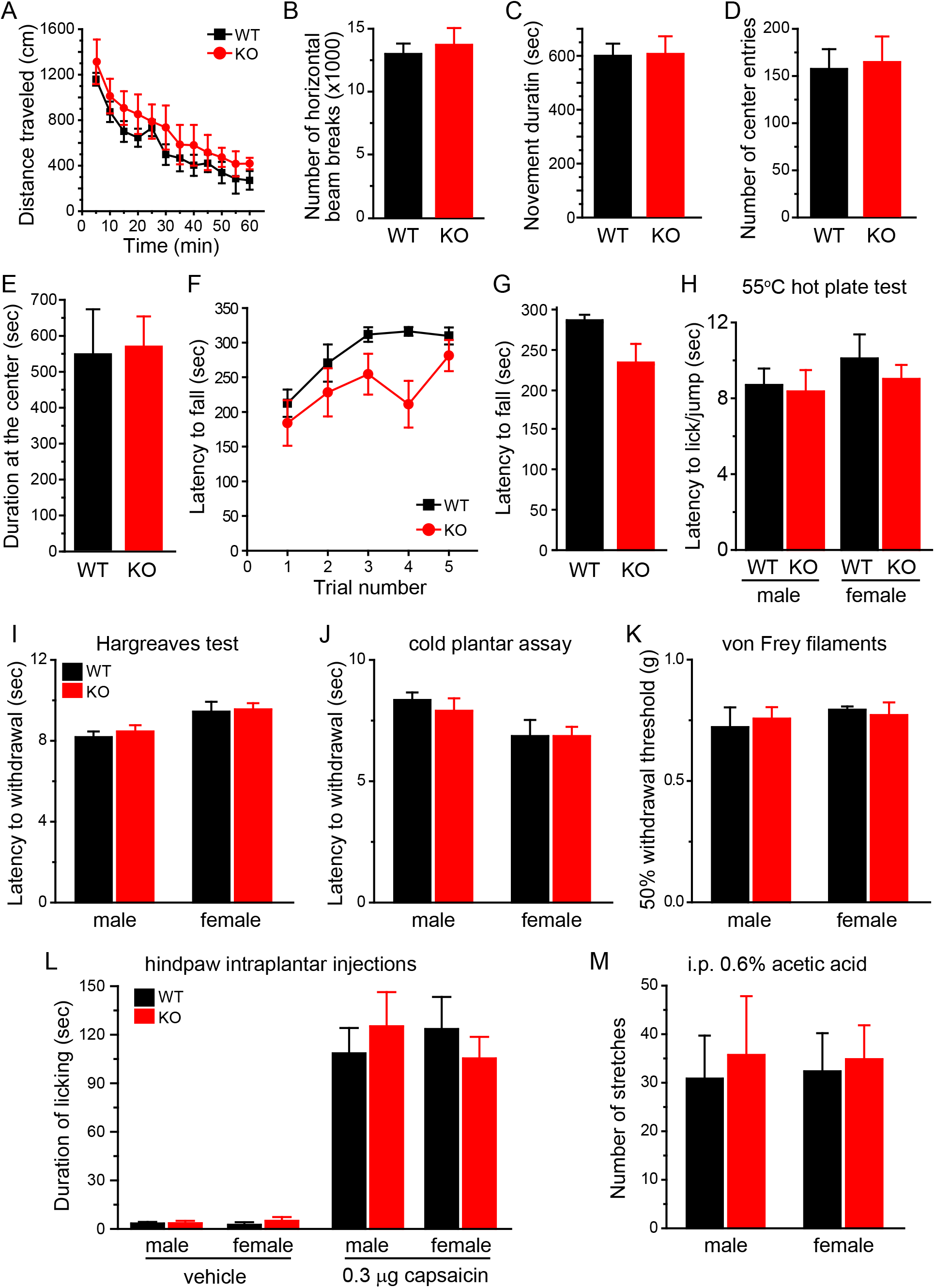
Both male and female TRESK KO mice show normal responses to stimuli on the hindpaw and in visceral tissues. (A) Distance traveled in the open field in 5-minute bins (n = 5 adult female and 5 adult male mice in each group). (B-E) Total number of horizontal beam breaks (B), total duration of movement (C), total number of entries into the center square (D) and total time spent in the center square (E) during the 60-minute testing period (same mice as in A). (F) Latency to fall from the accelerating rotarod in 5 consecutive trials (n = 5 adult female and 5 adult male mice in each group). (G) The latency to fall from the accelerating rotarod was averaged from the 5 trials in individual mice (same mice as in F). (H) Licking/jump latency on the 55°C hot plate (n = 5–6 mice in each group). (I) Withdrawal latency to radiant heat stimuli on the hindpaw (n = 5–6 mice in each group). (J) Withdrawal latency to cold stimuli on the hindpaw (n = 5–12 mice in each group). (K) Withdrawal threshold of a mechanical stimulus on the hindpaw (von Frey hair; n = 5–6 mice in each group). The results were verified by another experimenter in a separate cohort of WT and KO littermates. (L) Duration of licking the hindpaw after intraplantar injection of 10 µl vehicle (saline with 1% DMSO) or 0.3 µg capsaicin (n = 5–6 mice in each group). (M) The number of abdominal stretches produced by intraperitoneal injection of 0.6% acetic acid (n = 6–8 mice in each group).

First, we investigated whether genetic loss of TRESK affects the responses to noxious stimuli on the hindpaw and in visceral tissues. Previous studies reported that TRESK KO mice exhibited shorter response latency on a hot plate and a reduction of hindpaw mechanical threshold when housed individually (Castellanos et al., 2017; Chae et al., 2010). On the contrary, we found no difference between WT and KO mice in their responses to thermal and cooling stimuli to the hindpaw, regardless of sex (Figures 11H-J). The 50% withdrawal threshold to von Frey filaments on the hindpaw was also comparable between WT and KO mice (Figure 11K). This result was verified by another experimenter in a separate cohort of WT and KO littermates (data not shown). The discrepancy between the current and previous studies may arise from differences between genetic background (C57BL/6J versus C57BL/6N, plus the genetic drift in individual colonies), housing conditions (group housed to avoid social isolation stress versus individually housed) and diet composition. WT and KO mice also showed similar duration of licking in response to hindpaw injection of capsaicin (Figure 11L). Lastly, we tested the mice in a model of acute inflammatory visceral pain. The number of abdominal stretches evoked by intraperitoneal (i.p.) injections of dilute acetic acid did not differ in the WT and KO mice (Figure 11M). We conclude that genetic loss of TRESK preferentially enhances trigeminal pain across various modalities but does not alter the transmission and production of body pain, regardless of stimulus modality, target tissue and the sex of mice. This is consistent with the finding that ubiquitous loss of TRESK in PANs preferentially increases the intrinsic excitability and the responsiveness to capsaicin in TG neurons.

Secondly, we investigated whether endogenous TRESK activity contributes to trigeminal pain. We used an operant assay to assess the responses to noxious heat stimuli on facial skin. Mice needed to press their cheeks onto the Peltier bars set at various temperatures in the Orofacial Pain Assessment Device (OPAD) in order to lick the sweetened milk (the reward). In WT mice of either sex, both the total number of reward licking and the number of licks per facial contact (the L/F ratio) were greatly reduced when the bar temperature was increased from 33°C to 50°C (Figures 12A-B, black bars). TRESK KO mice responded similarly to WT mice at 33°C (Figures 12A-B, red bars), indicating they were not impaired in learning and performing the operant assay. However, the total number of licks and the L/F ratio at 50°C were significantly lower in both male and female KO mice relative to WT controls (Figures 12A-B), indicating that loss of TRESK increases the sensitivity and/or reduces the tolerance to noxious heat stimuli on facial skin. Interestingly, in female but not male TRESK KO mice, the number hot air (55°C)-evoked eye blinks was significantly higher than WT controls (Figure 12C).

**Figure 12.**
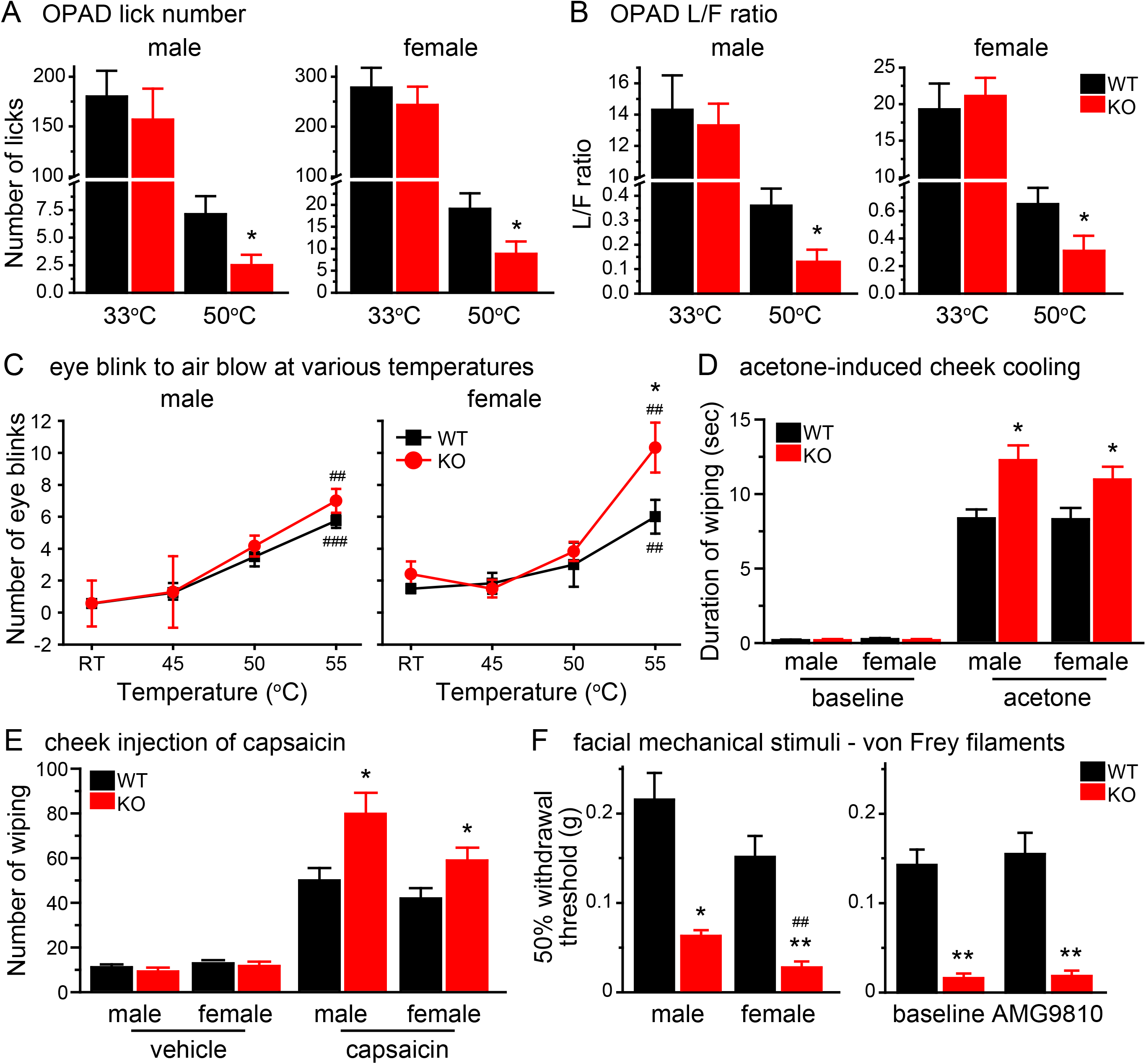
Loss of TRESK preferentially enhances trigeminal nociception. (A) Average number of reward licks with cheeks pressing the Peltier bars set at 33°C or 50°C during a 3-minute testing period in the OPAD assay (n = 7–11 mice in each group). *p < 0.05; two-tailed t-test between the corresponding WT and KO groups. (B) Average L/F ratio (the number of reward licks per cheek contact) during a 3-minute testing period in the OPAD assay (same mice as in A). (E) The number of eye blinks evoked by blowing 55°C air to the eye for 10 seconds (n = 6–8 mice in each group). (C) The number of eye blinks evoked by blowing air at various temperatures to the eye for 10 seconds (n = 6–8 mice in each group). *p < 0.05; two-tailed t-test between the corresponding WT and KO groups. ^##^p < 0.01; ^###^p < 0.001; one-way RM ANOVA with post hoc Bonferroni test, compared with the corresponding RT groups. (D) Time spent wiping the treated area after application of 15 µl acetone to the cheek (n = 10–12 mice in each group). Baseline activity was recorded for 1 min in individual mice before acetone application. *p < 0.05; two-tailed t-test between the corresponding WT and KO groups. (E) The number of forepaw wiping of the treated area after intradermal injection of 20 µl vehicle (saline with 1% DMSO) or capsaicin (1 µg in 20 µl vehicle) to the cheek (n = 7–13 mice in each group). *p < 0.05; two-tailed t-test between the corresponding WT and KO groups. (F) Left: the 50% withdrawal thresholds to punctate mechanical stimuli on forehead skin (n = 6– 8 mice in each group). Right: the 50% withdrawal thresholds in female mice (n = 8) before and after i.p. injection of TRPV1 antagonist AMG9810. *p < 0.05; **p < 0.01; Kruskal-Wallis ANOVA and Mann-Whitney U post hoc test with Bonferroni correction, between the corresponding WT and KO groups; ^##^p < 0.01, between male and female KO groups.

To test cold sensitivity, we reduced the facial skin temperature by applying acetone to the mouse cheek. The duration of wiping the treated area was significantly prolonged in KO mice than in WT (Figure 12D), indicating a hyper-responsiveness to acetone evaporation-induced cooling. To measure facial responses to chemical stimuli, we injected capsaicin intradermally into the mouse cheek and found that the number of cheek wiping was significantly increased in both male and female KO mice compared with WT controls (Figure 12E). Regarding facial mechanical sensitivity, we measured the threshold for evoking a withdrawal reflex to the application of von Frey filaments on mouse forehead skin. The 50% withdrawal threshold of TRESK KO mice was significantly lower than that of WT mice (Figure 12F, left). Notably, the withdrawal threshold of female KO mice was even lower than that of male KO, although WT mice did not show sex difference in mechanical thresholds (p = 0.32, Figure 12F, left). TRPV1 activation has been shown to contribute to mechanical allodynia in disease states (Shepherd et al., 2018; Yin et al., 2013). However, TRPV1 antagonist AMG9810 did not alter the facial mechanical withdrawal thresholds in WT or KO mice (Figure 12F, right), indicating that loss of TRESK is sufficient to increase the sensitivity to facial mechanical stimuli. Together, these results indicate that endogenous TRESK activity inhibits the transmission and production of facial pain evoked by thermal, cold, chemical and punctate mechanical stimuli.

### Dural afferent neurons from TRESK KO mice show higher intrinsic excitability and capsaicin responsiveness

Does genetic loss of TRESK affect the excitability of dural afferent neurons, the primary sensory neurons in the trigeminovascular pathway subserving headache? We used Fluoro-Gold (FG) to retrogradely label dural afferent neurons in adult mice and found that the size distribution of FG-labeled (FG^+^) dural afferent neurons was similar between WT and TRESK KO mice (Figure 13H). First, we compared the intrinsic excitability of WT and KO dural afferent neurons. Compared with WT small IB4^−^ dural afferent neurons, the mean rheobase was 50% lower in KO neurons and the average R_in_ was significantly higher (Figure 13A and Table 3). The number of APs evoked by depolarizing current injections was significantly increased in KO neurons relative to the WT group (Figure 13B). On the contrary, the rheobase, R_in_ and spike frequency were similar in small IB4^+^ dural afferent neurons from WT and KO mice (Figures 13A, C). The values of V_rest_, AP threshold, amplitude, half-width, and AHP amplitude were all comparable between WT and KO FG^+^ neurons (Table 3).

**Figure 13.**
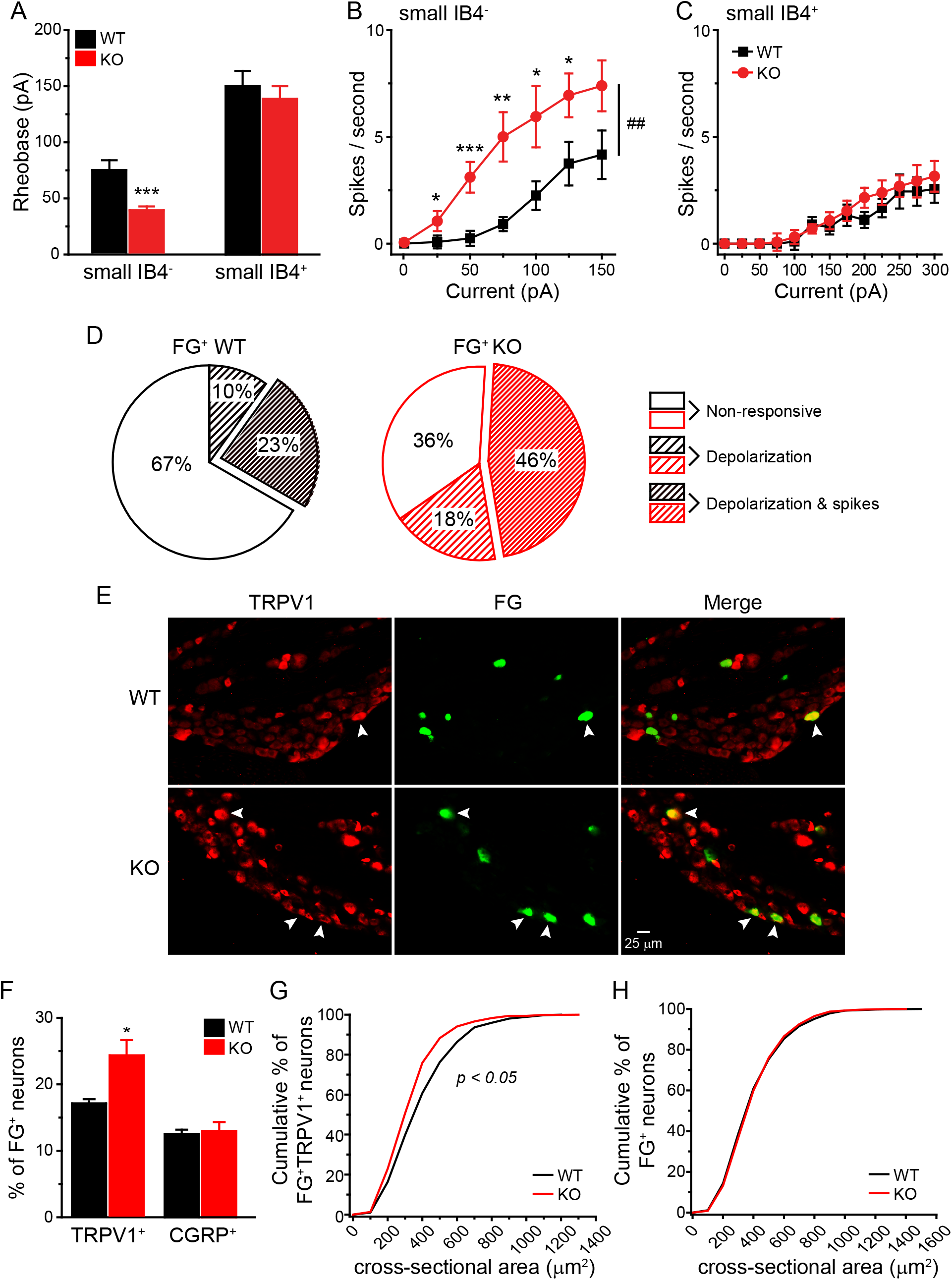
Dural afferent neurons from TRESK KO mice show higher intrinsic excitability and capsaicin responsiveness. (A) Mean rheobase of FG^+^ small IB4^−^ and IB4^+^ dural afferent neurons from WT and KO mice (n = 13–20 neurons in each group, see Table 3 for details of the intrinsic properties of dural afferent neurons). ***p < 0.001; two-tailed t-test between the corresponding WT and KO groups. (B-C) Input-output plots of the spike frequency in response to incremental depolarizing current injections in small IB4^−^ (B) and small IB4^+^ (C) dural afferent neurons from WT and KO mice (same neurons as in A). ^##^p < 0.01; two-way RM ANOVA; **p < 0.01 between the corresponding WT and KO groups. (D) The percentage of WT and KO dural afferent neurons that exhibit depolarization, burst of spikes or no response to 100 nM capsaicin (n = 30 WT and 28 KO neurons). p < 0.01; χ^2^ test between WT and TRESK KO groups. (E) Representative images of WT and KO TG sections that contain FG^+^ dural afferent neurons and TRPV1^+^ neurons. Arrowheads indicate neurons that are both FG^+^ and TRPV1^+^. (F) The abundance of TRPV1^+^ and CGRP^+^ dural afferent neurons in WT and KO mice (n = 3–4 mice in each group; on average, 708 FG^+^ neurons/mouse were counted). *p < 0.05, two-tailed t-test between the corresponding WT and TRESK KO groups. (G) Cumulative distributions of the cross-sectional areas of WT and KO FG^+^TRPV1^+^ dural afferent neurons (n = 387 and 581 neurons pooled from 4 WT and 4 KO mice, respectively, same mice as in F). p < 0.05; Mann–Whitney U test. (H) Cumulative distributions of the cross-sectional areas of FG^+^ dural afferent neurons in wild-type and TRESK KO mice (n = 3627 and 3332 FG^+^ neurons pooled from 4 WT and 4 KO mice, respectively, same mice as in F).

**Table 3.**
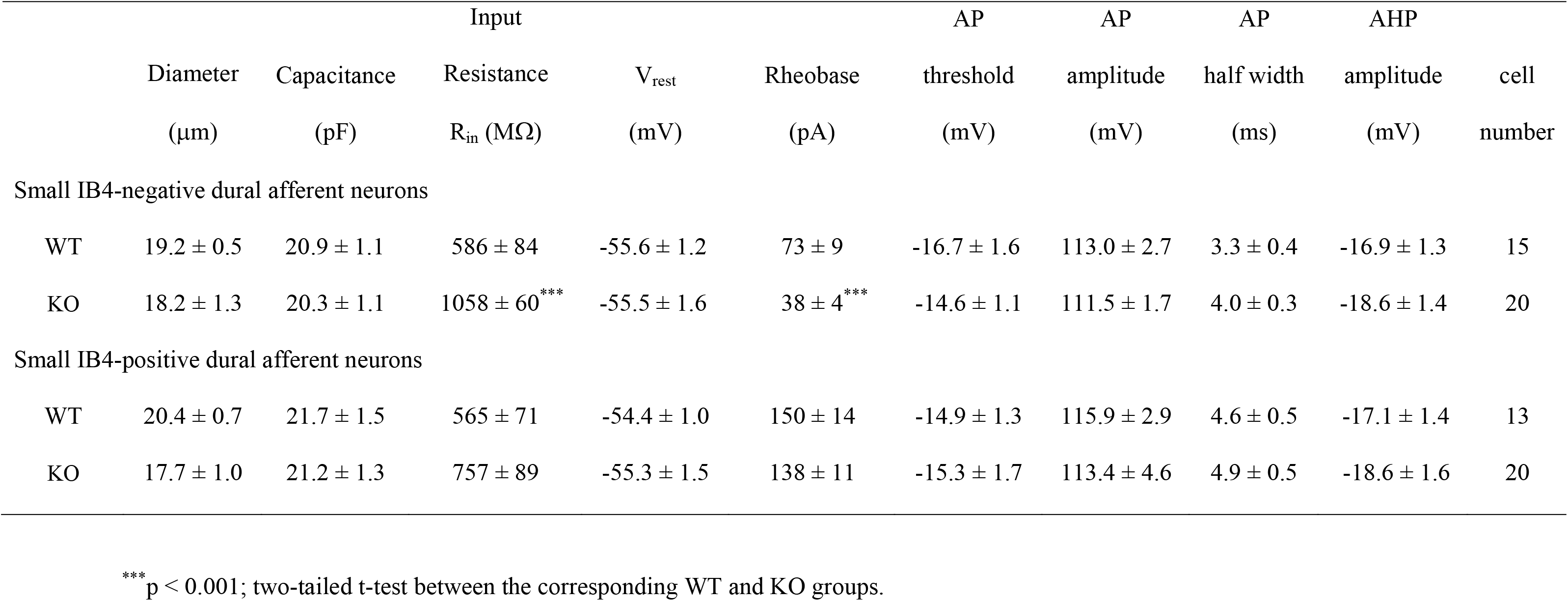
Intrinsic properties of FG-labeled dural afferent neurons from adult WT and TRESK KO mice.

Secondly, we recorded the responses of WT and KO dural afferent neurons to capsaicin under current-clamp. Bath application of 100 nM capsaicin evoked membrane depolarization and, in some cases, multiple spikes in 33% (10 of 30) WT neurons (Figure 13D black pie). Conversely, 46% (13 of 28) KO dural afferent neurons exhibited capsaicin-induced multiple spikes and an additional 18% (5 of 28) were depolarized by capsaicin (Figure 13D red pie). We also compared the expression of TRPV1 and CGRP between WT and KO dural afferent neurons. Consistent with a previous report (Huang et al., 2012), the abundance of TRPV1^+^ and CGRP^+^ neurons were lower in FG^+^ dural afferent neurons than in the total TG population in WT mice (Figures 8C, 13F). The percentage of TRPV1^+^ dural afferent neurons was significantly higher in KO mice than that in WT controls (Figures 13E, F), a more than 40% increase on average. The soma sizes of TRPV1^+^ dural afferent neurons in TRESK KO mice were slightly smaller than those in WT mice (Figure 13G). Conversely, the abundance of CGRP^+^ dural afferent neurons was comparable between WT and KO mice (Figure 13F). We conclude that loss of TRESK activity results in hyper-excitation of small IB4^−^ dural afferent neurons as well as an expansion of TRPV1 channel expression in dural afferents.

### TRESK KO mice are hyper-responsive to stimuli that generate headache-related behavior

Here, we investigated whether endogenous TRESK activity regulates the generation of headache-related behaviors in mice (Huang et al., 2016). After recovery from craniectomy for 7 days, both WT and TRESK KO mice exhibited some spontaneous forepaw wiping and hindpaw scratching behavior within the trigeminal V1 dermatome during a 2-hour observation period in the home cage (Figures 14A-D, baseline groups). Dural application of 20 µl vehicle did not increase V1-directed behavior above the basal level in either WT or KO mice (Figures 14A-D, vehicle groups). In WT mice, dural application of 20 µl IScap, which contained capsaicin and a mixture of inflammatory mediators, elicited more robust V1-directed wiping and scratching than vehicle treatment (Figures 14A-D, black bars). Our previous study suggests that dural IScap-induced behavior are mechanistically related to the ongoing headache in humans, as they can be reduced to the control level by pretreatment of mice with anti-migraine drugs (Huang et al., 2016). Both the number (Figures 14A-B) and the duration (Figures 14C-D) of V1-directed behaviors were significantly higher in TRESK KO mice than in WT mice regardless of sex, indicating that loss of TRESK renders both male and female mice hyper-responsive to stimuli that evoke headache-related behaviors.

**Figure 14.**
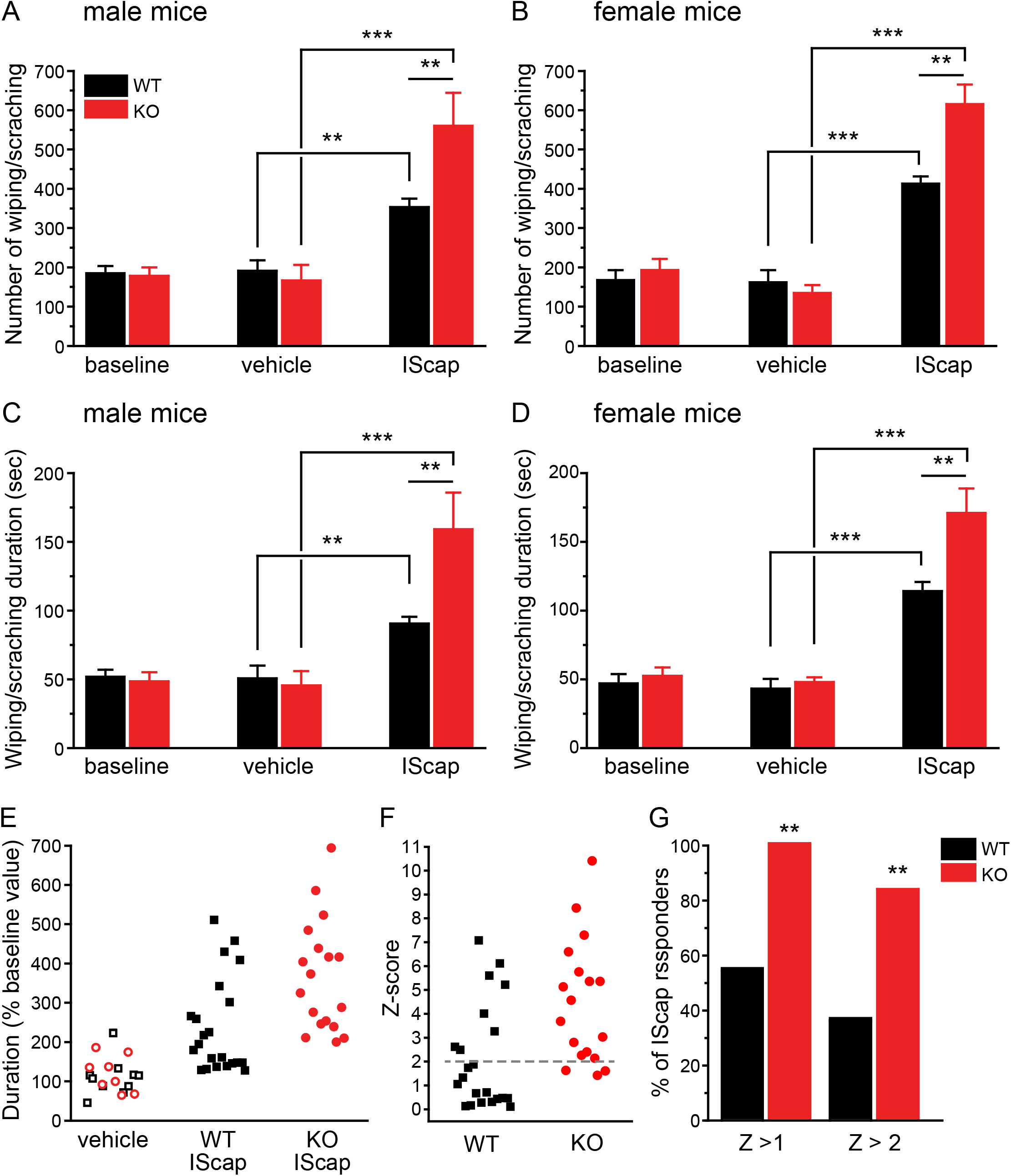
TRESK KO mice are hyper-responsive to dural application of IScap. (A) The number of V1-directed forepaw wiping and hindpaw scratching within the 2-hour recording period in male mice. Baseline behaviors were recorded in all mice 7 days post-surgery. The next day, mice received dural application of vehicle (n = 5 WT and 4 KO mice) or IScap (n = 10 WT and 8 KO mice). The behaviors were recorded 0.5 – 2.5 hours after the dural application. **p < 0.01; ***p < 0.001; two-way ANOVA with post hoc Bonferroni test. (B) The number of V1-directed behaviors in female mice (n = 5 WT and 4 KO mice in the vehicle groups; n = 12 WT and 10 KO mice in the IScap groups). (C-D) The duration of V1-directed forepaw wiping and hindpaw scratching within the 2-hour recording period in male (C) and female (D) mice (same mice as in A and B). (E) Scatter plots of the duration of V1-directed behavior (normalized to the baseline value in individual mice) in response to dural application of vehicle or IScap. Data from the male and female mice are combined. Data from vehicle-treated mice are pooled together (open black square and red circle indicate WT and KO mice, respectively). (F) The Z score of individual IScap-treated mice, calculated as Z = (value – mean) / SD. Value is the normalized duration of individual IScap-treated mice. Mean and SD are calculated from the normalized duration of all vehicle-treated mice. (G) The percentage of WT and KO mice that respond to dural application of IScap, using Z > 1 and Z > 2 (> mean duration + 1*SD or + 2*SD of vehicle-treated mice) as the threshold respectively. **p < 0.01, Fisher’s exact test between the corresponding WT and KO groups.

We went on to ask whether loss of TRESK alters the likelihood to manifest headache-related behaviors in mice. To increase statistical power, we combined data from male and female mice and normalized the duration of vehicle- or IScap-induced behavior to the baseline values in individual mice (Figure 14E). Since WT and KO mice responded similarly to vehicle application, we used all vehicle-treated mice as the reference population to calculate the Z score (the number of standard deviations [SDs] from the mean of all vehicle-treated mice) of each IScap-treated mice (Figure 14F). When we used z > 2 as the threshold, only less than 40% (8 of 22) of WT mice met the criteria as IScap responders; whereas more than 80% (15 of 18) TRESK KO mice were responders (Figure 14G). Using a less stringent criteria (Z > 1), all KO mice responded to dural IScap but less than 60% of WT mice could be classified as responders (Figures 14G). Collectively, these data suggest that loss of TRESK increases both the likelihood and the magnitude of headache-related behavior in response to IScap-evoked activation of the trigeminovascular pathway.

## Discussion

Although mutations of TRESK channels have been reported in migraine patients, whether TRESK activity controls the generation of trigeminal pain, especially headache, is still debated. The lack of specific blocker presents a major roadblock for deciphering the contribution of TRESK to sensory processing. To this end, we have used WT and TRESK KO mice to systematically investigate the impact of genetic loss of TRESK on trigeminal and body pain transmission.

### Cell type-specific effect of TRESK dysfunction on intrinsic excitability

Consistent with previous work (Dobler et al., 2007; Guo & Cao, 2014; Kollert et al., 2015; Lafreniere et al., 2010; Yoo et al., 2009), we found that TRESK-ir and lamotrigine-sensitive TRESK currents are present in every WT TG and DRG neuron. Loss of TRESK results in a significant reduction of total persistent outward current in all TG neurons. In small IB4^−^ TG neurons that express CGRP and/or TRPM8, this leads to a significant increase in R_in_, a decrease of rheobase, and an increase in spike frequency in response to depolarizing current injections, indicating that endogenous TRESK is activated around V_rest_ and during the weak depolarizations prior to AP initiation to reduce the membrane resistance and increase outward currents, thereby serving as a brake to oppose membrane depolarization and limit AP firing. Of note, TRESK is the only K_2P_ channel that can be activated by intracellular Ca^2+^ (Enyedi & Czirjak, 2015). In response to noxious stimuli, TRESK activity can be enhanced by the elevation of intracellular Ca^2+^, further counterbalancing the generator potential provided by excitatory inputs. In addition, Ca^2+^ influx through voltage-gated Ca^2+^ channel during AP repolarization also increases endogenous TRESK current, thereby reducing neuron’s ability to generate multiple APs.

Importantly, we observed spontaneous APs at V_rest_ in ∼10% of the small IB4^−^ TG neurons from TRESK KO mice but not in any of the WT TG neurons. What distinguishes these neurons from those without spontaneous APs was the lower AP threshold, suggesting that endogenous TRESK activity prevents spontaneous APs through delay of the activation of voltage-gated Na^+^ channels in some WT small IB4^−^ TG neurons. Numerous studies indicate that ongoing activity of PANs maintains the ongoing (“spontaneous”) pain under tissue- and nerve-injury conditions (Baron et al., 2013; Gracely et al., 1992; Haroutounian et al., 2014). Our findings may shed light on one of the mechanisms through which ongoing activity is generated in C-fibers, especially the small IB4^−^ TG neurons under pathological conditions.

Despite a similar increase in R_in_, small IB4^+^ TG neurons from TRESK KO mice only exhibited a 30% reduction of rheobase, with no changes in spike frequency and no occurrence of spontaneous APs at V_rest_, indicating a limited role of endogenous TRESK in regulating the excitability of this TG subpopulation. This is in line with earlier studies identifying another K_2P_ channel TREK2 as well as Ca^2+^- and Na^+^-modulated K^+^ channels as key regulators of the V_rest_ and intrinsic excitability of small IB4^+^ PANs (Acosta et al., 2014; Martinez-Espinosa et al., 2015; Zhang et al., 2010). Interestingly, although functional TRESK channels are clearly present in medium-sized TG neurons, there is little TRESK activity when the membrane potential is below −35 mV. This may explain why loss of TRESK did not affect the passive membrane properties or the excitability of medium-sized TG neurons at all. Whether the gating of TRESK channel is different in medium- and small-diameter TG neurons requires further investigation.

In contrast to what we observed in TG neurons, loss of TRESK did not reduce total persistent outward current or alter the intrinsic excitability of DRG neurons. Upregulation of other K^+^ channels that are active at V_rest_ (e.g. other K_2P_ family of channels) likely compensates for the loss of endogenous TRESK. Further study is needed to determine which channel(s) are upregulated in response to genetic loss of TRESK in DRG, how long it takes to restore background K^+^ current and, importantly, why this adaptive change does not occur in TG neurons. A comparison of the transcriptomes from mouse TG and DRG neurons reveals that, despite being overwhelmingly similar, 15 members of the Hox family of transcription factors are exclusively expressed in adult DRG neurons (Lopes et al., 2017). This provides a possible molecular basis for the differential adaptive changes of DRG and TG neurons in response to the loss of TRESK.

Collectively, our present data delineated a cell type-specific role of TRESK channels in controlling TG neuronal excitability. Despite the ubiquitous presence of TRESK activity in all PANs, genetic loss of TRESK preferentially increases the excitability of small IB4^−^ TG nociceptors expressing CGRP and/or TRPM8. Notably, a moderate reduction of TRESK and total persistent outward currents had no effect on the intrinsic excitability of heterozygous TG neurons, indicating that 50% of endogenous TRESK activity is sufficient to maintain normal intrinsic excitability and to prevent spontaneous APs. A more profound reduction of TRESK activity would be required to cause prolonged hyper-excitation of subpopulations of TG neurons during migraine and other chronic trigeminal pain conditions. In a recent study, transient reduction of TRESK current with a dominant-negative subunit does not alter the excitability of small TG neurons (Royal et al., 2018). However, it is not clear to what extent the TRESK and total persistent outward currents are affected by the mutant subunit. More importantly, including all small TG neurons into one group would likely mask the effects of the mutant subunit on the excitability of subpopulations of TG neurons, as indicated by our results.

### Novel function of TRESK in regulating TRPV1 expression

Previous studies of K_2P_ channel functions have focused on their contributions to neuronal excitability. Loss of other K_2P_ channels such as TRAAK, TREK1, TREK2 and/or TASK3 increases the percentage of DRG neurons that respond to capsaicin, menthol as well as temperature changes through attenuating the brake for excitability (Morenilla-Palao et al., 2014; Noel et al., 2009; Pereira et al., 2014). In the present study, loss of TRESK nearly doubled the number of Cap^+^ TG neurons as the result of an increase in the number of neurons that express TRPV1 mRNA and protein. Our findings revealed for the first time a function of TRESK beyond its role in regulating neuronal excitability.

The expression of TRPV1 occurs in a wide range of PANs in early development and gradually become more restricted to a subset of PANs in adult mice (Cavanaugh et al., 2011; Luo et al., 2007; Mishra et al., 2011; Takashima et al., 2010). Although the developmental onset of TRESK expression in TG neurons is not well defined, the density of TRESK current in neonatal TG neurons is already comparable to that of adult TG neurons (Guo & Cao, 2014); whereas the refinement of TRPV1 expression does not complete till postnatal day 15 (Cavanaugh et al., 2011). Loss of TRESK may perturb this process, resulting in TRPV1 expression in a larger fraction of TG neurons. Alternatively, the expansion of TRPV1 expression may be activity-dependent, resulting from prolonged hyper-excitation of small IB4^−^ TG neurons without TRESK. Future research is warranted to investigate these non-mutually exclusive possibilities. Whether TRESK dysfunction contributes to the upregulation of TRPV1 in adult PANs under some chronic condition also merits further study. All in all, our experiments provide a starting point for further elucidating the potential role of TRESK channels in regulating gene expression in addition to serving as the brake for AP generation.

The present results also firmly establish that the small IB4^−^ TG neurons are the PANs that are most vulnerable to TRESK dysfunction, undergoing maladaptive changes to increase the intrinsic excitability and to express more sensor proteins to detect noxious stimuli. It is well recognized that TRPV1 can be activated by a variety of endogenous and exogenous stimuli (Caterina & Julius, 2001). The expansion of TRPV1^+^ neurons mainly occurs in KO TG neurons that express NF200, CGRP and/or TRPM8, suggesting that some mechano- or cold-sensitive TG neurons are converted to polymodal nociceptors in the absence of TRESK. This likely would enhance the ability of individual PANs in detection and integration of noxious stimuli of various modalities in both normal and chronic pain states.

### Implication of TRESK dysfunction for trigeminal pain, especially migraine headache

Both our *in vitro* experiments and the behavioral tests reveal no difference between WT and TRESK KO mice in their responses to noxious stimuli on the hindpaw and in viscera. This does not imply that TRESK activity is irrelevant to body and/or visceral pain in adults, but rather it supports the view that DRG neurons may in some way compensate for the genetic loss of TRESK. Earlier studies have provided clear evidence that reduction of TRESK activity in adult mice increases the sensitivity to mechanical stimuli on the hindpaw (Bautista et al., 2008; Lennertz et al., 2010; Tulleuda et al., 2011; Zhou et al., 2016). Future studies will be required to assess whether mature DRG neurons can restore the level of TRESK current long after the initial injury and whether this contributes to the resolution of pain.

We made several key *in vivo* observations that endogenous TRESK activity regulates trigeminal nociception across various modalities in both male and female mice. First, TRESK KO mice showed enhanced responses to noxious stimuli on facial skin but not on hindpaw skin, suggesting that loss of TRESK differentially affects the excitability of TG and DRG neurons that innervate cutaneous tissues. Thus, it seems unlikely that the selective increase in TG neuronal excitability in KO mice results from the unique target tissues (e.g. dura, eyes and tooth pulps) that some TG neurons innervate. Secondly, both the hyper-excitation of small IB4^−^ TG neurons, especially the CGRP^+^ neurons, and the increase in TRPV1 channel expression account for the hyper-sensitivity of TRESK KO mice to facial thermal and chemical stimuli (McCoy et al., 2013). It is well established that injury induces thermo-hypersensitivity through enhancing TRPV1 activity and/or expression (Caterina & Julius, 2001). In addition, injury-induced decrease of TRESK expression in PANs may increase the intrinsic excitability, thereby allowing heat-sensing neurons to respond to warm temperature (Yarmolinsky et al., 2016). Thirdly, loss of TRESK increases facial mechanical sensitivity independent of TRPV1 activation. Under normal condition, punctate mechanical sensation in mice is mediated by the non-peptidergic PANs that do not express TRPV1 (Cavanaugh et al., 2011). It is likely that a mild reduction of rheobase in small IB4^+^ TG neurons is sufficient to increase facial mechanical sensitivity in KO mice. Notably, loss of TRESK results in a more profound reduction of withdrawal threshold in female mice, and hot air-evoked eye blink was enhanced in female but not male KO mice, suggesting that TRESK dysfunction affect some aspects of trigeminal pain processing more severely in females than males. Lastly, the response to acetone-induced cooling of facial skin is more robust in TRESK KO mice, likely resulting from the hyper-excitation of TRPM8-expressing TG neurons. This indicates that endogenous TRESK, probably along with other K^+^ channels including TREK1, TREK2 and TASK3 (Madrid et al., 2009; Morenilla-Palao et al., 2014; Noel et al., 2009; Pereira et al., 2014; Vetter et al., 2013; Viana et al., 2002), acts as an excitability brake in TRPM8-expressing neurons to modulate the sensitivity to cold stimuli. Future experiments are necessary to determine whether the decrease of TRESK expression contributes to injury-induced cold-hypersensitivity by weakening the excitability brake.

Most importantly, we found that loss of TRESK selectively increases the intrinsic excitability and TRPV1 expression in small IB4^−^ dural afferent neurons. This was sufficient to increase the percentage of mice responsive to the dural application of IScap as well as to increase the magnitude of headache-related behaviors. These results indicate that the endogenous TRESK activity in small IB4^−^ dural afferent neurons negatively regulates the activation of the trigeminovascular pathway, thereby preventing the initiation of the headache episodes. Of note, many of the small IB4^−^ dural afferent neurons express CGRP and/or its receptors (Burstein et al., 2015; Ho et al., 2010). In response to migraine triggers, dural afferent neurons release CGRP and many other neurotransmitters/modulators in both meninges and medullary dorsal horn. CGRP can in turn activate more dural afferent neurons by signaling downstream of CGRP receptors (Burstein et al., 2015; Ho et al., 2010). In future studies, it will be necessary to investigate whether exposure to common migraine triggers causes a substantial reduction of endogenous TRESK activity in small IB4^−^ dural afferent neurons from WT mice. If this is the case, hyper-excitation of these neurons will not only enhance the release of CGRP but also amplify the effects of CGRP receptor signaling, thereby strengthening this local feedforward loop and leading to the initiation and maintenance of headache in patients with common forms of migraine.

TRESK mRNA is highly expressed in human TG neurons and dominant-negative TRESK mutations have been identified in migraine patients (Andres-Enguix et al., 2012; Flegel et al., 2015; Lafreniere et al., 2010; LaPaglia et al., 2018; Rainero et al., 2014). However, the causal relationship between TRESK function and migraine susceptibility has not been established. A recent study suggests that dysfunction of TREK1/2 channels, and not of TRESK alone, contributes to the increase TG neuronal excitability and alter the pain processing in migraine patients with frameshift TRESK mutations (Royal et al., 2018). We also found that a 50% reduction of endogenous TRESK activity does not alter the intrinsic excitability of TG neurons. That said, what we have observed in TRESK KO mice strongly suggest that a profound reduction of endogenous TRESK activity will significantly increase the excitability of small IB4^−^ dural afferent neurons and enhance the activation of the trigeminovascular pathway, leading to a higher susceptibility to headache episodes. Future research is warranted to test whether migraine triggers can profoundly reduce TRESK activity in dural afferent neurons, and whether this can precipitate headache episodes in common forms of migraine.

In humans, TRESK mutations are associated with migraine but not body pain or visceral pain. Similarly, TRESK KO mice exhibit behaviors indicative of enhanced trigeminal pain, including headache, but normal body pain and visceral pain; whereas genetic loss of TREK1 or TREK2 lead to hyper-sensitivity to noxious thermal, mechanical and chemical stimuli on the hindpaw, indicating enhance body pain responses (Alloui et al., 2006; Pereira et al., 2014). Under our housing and experimental conditions, TRESK KO mice better recapitulate the symptoms caused by human TRESK mutations than TREK1/2 KO mice. Admittedly, our findings using the global TRESK KO mice do not exclude a role for TRESK outside PANs. Future studies using tissue-specific KO mice will unequivocally demonstrate the contributions of PAN TRESK activity to chronic pain, especially migraine headache.

In conclusion, our study highlights some exquisite differences between TG and DRG neurons in response to ion channel dysfunction. We provide evidence that ubiquitous loss of TRESK during development preferentially increases the intrinsic excitability of TG nociceptors expressing CGRP or TRPM8 as well as expands the expression of TRPV1 proteins in small IB4^−^ TG neurons. Consequently, TRESK dysfunction selectively enhances trigeminal pain and, in particular, the generation of headache. Our findings support a causal relationship between TRESK function and migraine susceptibility and establish a foundation for further elucidating the unique molecular and cellular basis of trigeminal pain, especially migraine headache.

## Materials and Methods

### Mice

All procedures were carried out in strict accordance with the recommendations in the Guide for the Care and Use of Laboratory Animals of the National Institutes of Health and the guidelines of the Institutional Animal Care and Use Committee at Washington University in St. Louis. To avoid social isolation stress, all mice (except for the ones that underwent craniectomy) were group housed (2-5 per cage, same sex) in the animal facility of Washington University in St Louis on a 12-hour light–dark cycle with constant temperature (23-24°C), humidity (45-50%), and food and water *ad libitum*. All mice were maintained on C57BL/6J background (backcrossed for at least 7 generations). Young adult mice (5-7 weeks old) or neonatal mice (postnatal day 1-2) were used in the electrophysiology and Ca^2+^ imaging experiments. Adult male and female littermates (8-20 weeks old) were used in the behavioral tests and immunohistochemistry (IHC) experiments.

WT and TRESK KO mice were generated by crossing heterozygous breeders (Kcnk18^tm1(KOMP)Vlcg^, KOMP repository). We also crossed heterozygous TRESK breeders with heterozygous TRPM8^EGFP^ and VGLUT3^EGFP^ mice. The double heterozygous breeders were then crossed with TRESK heterozygotes to generate WT/KO_TRPM8^EGFP^ and WT/KO_VGLUT3^EGFP^ mice, respectively. Genotypes were determined by PCR of tail DNA as described (Dhaka et al., 2008; Seal et al., 2009; Valenzuela et al., 2003). For TRESK allele genotyping, the WT allele was amplified with primers TUF (5’-GAGGAGAACCCTGAGTTGAAGAAG-3’) and TUR (5’-GCACCTCCGAGGCAGTAAC-3’), producing a 103-bp fragment. The targeted allele was amplified with primers NeoInF (5’-TTCGGCTATGACTGGGCACAACAG-3’) and NeoInR (5’-TACTTTCTCGGCAGGAGCAAGGTG-3’), producing a 282-bp fragment. The PCR conditions were 96°C for 30 seconds (s), 50°C for 30 s and 72°C for 30 s for 40 cycles.

### Primary culture of mouse TG and DRG neurons

TG and lumbar DRG (L3-L5) tissues were collected from 5-6 weeks old WT and TRESK KO mice of either sex and were treated with 2.5 mg/ml collagenase IV followed by 2.5 mg/ml trypsin at 37°C for 15 minutes (min), respectively. Cells were dissociated by triturating with fire-polished glass pipettes and were centrifuged through a 15% BSA gradient to remove debris and non-neuronal cells. Neurons were resuspended in MEM-based culture medium containing 5% fetal bovine serum, 25 ng/ml NGF, 10 ng/ml GDNF and were seeded on Matrigel-coated coverslips. Ca^2+^ imaging and electrophysiology recordings were performed in neurons 2-4 DIV. For Ca^2+^ imaging experiments, the BSA gradient was omitted to increase the yield of neurons. In some cases, neurons were cultured in the absence of NGF and GDNF and were used within 24 hours after plating. Each set of experiment contains neurons from at least 3 batches of culture.

### Ca^2+^ Imaging

Coverslips containing cultured TG or DRG neurons were incubated with HBSS/Hepes solution containing 2.5 µM fura-2 AM and 0.1% Pluronic F-127 (all from Molecular Probe) at 37°C for 30 min to load the ratiometric Ca^2+^ indicator. De-esterification of the dye was carried out by washing the coverslips 3 times with HBSS/Hepes solution and incubating the coverslips in HBSS/Hepes solution in the dark for an additional 30 min at 37°C. Neurons were used for Ca^2+^ imaging experiments within 3 hours after Fura-2 loading.

Coverslips with fura-2 loaded neurons were placed in a flow chamber mounted on a Nikon TE2000S inverted epifluorescent microscope and were perfused with room temperature (RT) Tyrode’s solution (1 ml/min) containing (in mM): 130 NaCl, 2 KCl, 2 CaCl_2_, 2 MgCl_2_, 25 Hepes, 30 glucose, pH 7.3 with NaOH, and 310 mosmol/kgH_2_O. A differential interference contrast (DIC) image of neurons in the field was captured to calculate soma diameters from cross-sectional areas off-line. Healthy neurons were chosen based on their morphology under DIC microscopy. In a pilot test, all selected neurons (> 100) exhibited robust Ca^2+^ influx in response to a depolarization stimulus (extracellular solution containing 50 mM KCl). Fura-2 was alternately excited by 340 and 380 nm light (Sutter Lambda LS) and the emission was detected at 510 ± 20 nm by a UV-transmitting 20x objective (N.A. 0.75) and a CoolSnapHQ2 camera (Photometrics). The frame capture period was 50 milliseconds (ms) at 1.5 s interval. SimplePCI software (Hamamatsu) was used for controlling and synchronizing the devices as well as image acquisition and analysis. Regions of interest (ROIs) encompassing individual neurons were defined *a priori*. The ratio of fluorescence excited by 340 nm divided by fluorescence excited by 380 nm (R_340/380_) was determined on a pixel-by-pixel basis and was averaged for each ROI. An additional background area was recorded in each field for off-line subtraction of background fluorescence.

To measure capsaicin-induced Ca^2+^ influx, Tyrode’s solutions containing 100 nM or 1 µM capsaicin were prepared from a 50 µM capsaicin stock solution (in 100% ethanol) before each experiment. After a 3 min baseline measurement in Tyrode’s solution, neurons were perfused with 100 nM capsaicin at 5 ml/min for 1 min followed by washing with Tyrode’s for 5 min and perfusion with 1 µM capsaicin for 1 min. To measure the responsiveness to multiple chemical stimuli in individual neurons, we used a protocol consisted of a 3 min baseline measurement followed by 1 min perfusion of 1 µM capsaicin, 5 min Tyrode’s wash, 1 min perfusion of 100 µM histamine or 100 µM menthol, 5 min of Tyrode’s wash and 1 min perfusion of 100 µM mustard oil (all from Sigma). At the end of each experiment, neurons were incubated with 3 µg/ml Fluorescein isothiocyanate (FITC)-conjugated IB4 (Sigma) for 5 min. The FITC fluorescence on soma membrane was detected after 10 min perfusion with Tyrode’s solution to wash off unbound IB4.

Peak responses were determined by calculating the absolute increase in R_340/380_ above baseline (the average R_340/380_ during the 3 min baseline measurement). A > 20% change from baseline was set as the threshold for a response. Since R_340/380_ is directly related to intracellular Ca^2+^ concentration (Grynkiewicz et al., 1985), we used the average fold change of peak R340/380 values ([peak_R340/380 – basal_R340/380] / basal_R340/380) to compare the magnitude of Ca^2+^ transients.

### Electrophysiology

Whole cell patch-clamp recordings were performed at RT with a MultiClamp 700B amplifier (Molecular Devices). The recording chamber was perfused with Tyrode’s solution (0.5 ml/min). The pipette solution contained the following (in mM): 130 K-gluconate, 7 KCl, 2 NaCl, 0.4 EGTA, 1 MgCl_2_, 4 ATP-Mg, 0.3 GTP-Na, 10 HEPES, 10 Tris-phosphocreatine, 10 U/ml creatine phosphokinase, pH 7.3 with KOH, and 290 mosmol/kgH_2_O. Recording pipettes had < 4.5 MΩ resistance. pClamp 10 (Molecular Devices) was used to acquire and analyze data. Cell capacitance and series resistance were constantly monitored throughout the recording. Data were analyzed with the Clampfit (Molecular Devices) and Origin (OriginLab) software.

Neurons were recorded between 2-4 DIV. Longer culture time did not alter the number or the thickness of the processes but significantly increased the length and the branches of individual processes. The processes of cultured neurons would contribute to the space-clamp error. We did not find significant differences in the size of persistent outward current or neuronal excitability between early (2 DIV) and late (4 DIV) recordings within individual experimental groups. Thus the space clamp issue was not exacerbated by the prolonged culture time.

#### Voltage-clamp experiments

Series resistance (< 15 MΩ, average 12 ± 1 MΩ) was compensated by 80%. Current traces were not leak subtracted. Signals were filtered at 2 kHz and digitized at 20 kHz. Total background K^+^ current and current through TRESK channels in neurons were recorded as described previously (Guo & Cao, 2014). Briefly, the extracellular solution contained 1 µM tetrodotoxin (TTX) to block TTX-sensitive Na^+^ current. To minimize other transient voltage-gaged Na^+^, K^+^ and Ca^2+^ currents, we held neurons at −60 mV and depolarized them to −25 mV for 150 ms, and then ramped the potential to −135 mV at 0.37 mV/ms every 10 s. The outward current at the end of the −25 mV depolarizing step was measured as the total background K^+^ current. To further dissect background K^+^ currents through TRESK channels, we bath-applied 30 µM lamotrigine (Sigma) while evoking whole cell currents using this pulse protocol (Guo & Cao, 2014).

#### Current-clamp experiments

Series resistance (< 15 MΩ) was not compensated. Signals were filtered at 10 kHz and digitized at 100 kHz. After whole cell access was established, membrane capacitance was determined with amplifier circuitry. The amplifier was then switched to current-clamp mode to measure V_rest_. R_in_ was calculated by measuring the membrane potential change in response to a 20 pA hyperpolarizing current injection from V_rest_. Neurons were excluded from analysis if the V_rest_ was higher than −40 mV or R_in_ was smaller than 200 MΩ.

To test neuronal excitability, neurons at V_rest_ were injected with 1 s depolarizing currents in 5 or 25 pA incremental steps. The rheobase was defined as the minimum amount of current required to elicit at least 1 AP. The first AP elicited using this paradigm was used to measure AP threshold (the membrane potential at which dV/dt exceeds 10 V/s), amplitude, and half-width. The AHP amplitude was measured from the single AP elicited by injecting a 1 ms depolarizing current in 200 pA incremental steps from the V_rest_.

DIC images of neurons were captured before the recording to calculate soma diameters from cross-sectional areas off-line. At the end of electrophysiological recording, neurons were stained with FITC-conjugated IB4 (3 µg/ml). The recording pipette remained attached to the neurons during IB4 staining and washing. The V_rest_, R_in_, capacitance, series resistance, and leak currents were not significantly altered during the process.

### RNA Extraction, Reverse Transcription and qPCR

We followed the procedures described in a previous study (C. Yang et al., 2011). Briefly, TG tissues were collected from naive adult mice and total RNA was isolated using the RNeasy Plus Mini Kit (Qiagen). Two micrograms of total RNA from each sample was used for first-strand cDNA synthesis using the Retroscript Kit (Applied Biosystems, ABI). The reverse transcriptase was omitted in the negative control groups for qPCR.

The qPCR reactions were performed with samples in triplicate on an ABI 7500 fast real-time PCR system using power SYBR green PCR master mix (ABI). The mouse TRPV1 cDNA was amplified with primers 5’-TTCCTGCAGAAGAGCAAGAAGC-3’ and 5’-CCCATTGTGCAGATTGAGCAT-3’ (Albers et al., 2006). The mouse β-actin cDNA was amplified with primers 5’-TGGAGAAGAGCTATGAGCTGCCTG-3’ and 5’-GTAGTTTCATGGATGCCACAGGAT-3’ (C. Yang et al., 2011). The PCR condition was 95°C for 15 s and 60°C for 30 s for 40 cycles. The specificity of qPCR was verified by dissociation-curve analysis in both experimental and negative control groups. Primer efficiency was validated as described (C. T. Yang et al., 2009). The TRPV1 and β-actin primers exhibit similar efficiencies (94% and 90%, respectively). The threshold cycle (Ct) values were measured from the point where the linear range of the amplification curve intersects the amplification detection threshold line. Abundance of TRPV1 mRNA in each sample was calculated from 2^−dCt^, with dCt = Ct(TRPV1) − Ct(β-actin). Ct(TRPV1) and Ct(β-actin) represented the mean values of the triplicate reactions. The fold change in TRPV1 mRNA expression in individual TGs was calculated with the 2^-ddCt^ method (Livak & Schmittgen, 2001), with ddCt = dCt (each TG) – mean dCt of WT TGs.

### Retrograde labeling of TG neurons innervating the dura

We used FG (2% in saline, Fluorochrome) to label the dural afferent neurons in WT and TRESK KO mice as described in a previous study (Huang et al., 2012). Briefly, adult mice were anesthetized with 3-4% isoflurane in an induction chamber until losing the righting reflex and were mounted on a Stoelting stereotaxic apparatus. Anesthesia was maintained by 1.5-2% isoflurane through a nose cone. Body temperature was maintained by placing mice on a 37°C circulating water warming pad. The eyes were covered by a small amount of eye drops to prevent the corneas from drying. A longitudinal skin incision was made to expose the cranium. The muscle and periosteal sheath were removed. Lidocaine hydrochloride jelly (2%) was applied on the skin and the skull for 5 min before the incision and the muscle/sheath removal to prevent the activation and/or sensitization of the primary afferent fibers. A craniectomy (∼2.5 mm diameter) was made with a surgical blade in the area overlying the superior sagittal sinus (SSS) between the bregma and lambda, leaving the underlying dura exposed but intact. To prevent spreading of the tracer to other peripheral sites, a sterile polypropylene ring was sealed to the skull surrounding the exposed dura using a mixture of dental cement powder (Stoelting 51459) and superglue adhesive. After waiting 5–10 min for the mixture to solidify, we applied 20 μl FG onto the exposed dura. Subsequently, a sterile polypropylene cap was secured over the ring with the dental cement/superglue mix to cover the exposed dura. The skin incision was closed with 5-0 suture. After recovery from anesthesia, mice were housed individually in the animal facility to allow the transportation of FG to the somata of dural afferent neurons in TG. For electrophysiology experiments, craniectomy and FG application was done in 6 weeks old mice and TG tissues were collected 7 days later for primary culture. For IHC experiments, mice were perfused 14 days after the dural application of FG to collect TG tissues.

### Tissue Preparation, IHC, and Image Analysis

Adult WT and TRESK KO mice were euthanized with i.p. injection of barbiturate (200 mg/kg) and were transcardiacally perfused with 0.1 M phosphate-buffered saline (PBS, pH 7.2) followed by 4% formaldehyde in 0.1 M phosphate buffer (PB, pH 7.2) for fixation. TGs and DRGs were dissected out, post-fixed for 4 hours, and protected overnight at 4°C in 0.1 M PB with 30% sucrose. The entire TG or DRG was then frozen in Optimal Cutting Temperature compound, sectioned at 15 µm in the transverse plane using a cryostat, collected on Superfrost Plus glass slides in sequence and stored at −20°C.

One in every 3 TG sections was processed for each IHC experiment. The sections were dried at RT, washed 3 times in 0.01 M PBS, and incubated in blocking buffer consisting 0.01 M PBS, 10% normal goat serum (NGS), and 0.3% triton X-100 for 1 hour at RT. Sections were then incubated overnight with primary antibodies diluted in blocking buffer in a humidity chamber at 4°C. After 6 washes (5 min each) in washing buffer containing 0.01 M PBS with 1% NGS and 0.3% triton and 1 hour incubation in blocking buffer, sections were incubated with secondary antibodies (1:1000 dilution in blocking buffer) at RT for 1 hour, washed 6 times with washing buffer and rinsed 3 times in 0.01 M PBS. Sections were then cover-slipped using Fluoromount-G Slide Mounting Medium (Electron Microscopy), sealed with nail polish, and stored at 4 °C.

To count total TG neurons, sections were stained with the rabbit anti-βIII tubulin antibody (Covance, 1:1000) and AlexaFluor 568-conjugated goat anti-rabbit secondary antibody. Images of the entire TG section were captured using an Olympus NanoZoomer Whole-Slide Imaging System at the Alafi neuroimaging core facility at Washington University Medical School. All neurons on each section were counted and the total number of βIII tubulin-positive neurons was multiplied by 3 to obtain the total number of TG neurons for each mouse (Golden et al., 2010).

Other primary antibodies used in IHC were: rabbit anti-Iba1 (Wako, 1:1000), rabbit anti-TRPV1 (Neuromics, 1:1000), guinea pig anti-CGRP (Peninsula laboratories, 1:1000), mouse anti-NF200 (Sigma, 1:1000), chicken anti-EGFP (AVES Lab, 1:1000); and mouse anti-TRESK (1:1000, (Guo & Cao, 2014; Liu et al., 2013)). AlexaFluor 568- and 488-conjugated secondary antibodies (Invitrogen) were used at 1:1000 dilution.

High-power images of the TG sections were captured through a 20x objective on a Nikon TE2000S inverted epifluorescence microscope equipped with a CoolSnapHQ2 camera (Photometrics). Cross-sectional somatic area was measured using the SimplePCI software (Hamamatsu). Representative images were adjusted for contrast and brightness using the same parameter within individual experiments. No other manipulations were made to the images.

### Behavioral Tests

WT and TRESK KO mice were generated by crossing heterozygous breeders on C57BL/6J background. Adult male and female littermates (8-20 weeks old) were used in the behavioral tests. The experimenters were blinded to the genotype of mice and the treatments mice received.

### General motor function tests

#### Open field test

Mice were habituated in the testing room for 1 hour in their home cages and then tested one at a time. Each mouse was placed in the center of the illuminated, sound-attenuated VersaMax Open Field box (42 × 42 cm, AccuScan Instruments) for 1 hour while the experimenter left the room. Horizontal movement and center entries (14 × 14 cm) were recorded and analyzed by the VersaMax software.

#### Rotarod test

Mice were habituated in the testing room for 1 hour in their home cages and then underwent 5 training sessions on the Rotarod (Ugo Basile, Model 7650) at 4 revolutions/min (rpm) for 5 min. Mice that stayed on the rod for at least 2 min / session were tested on the accelerating rod (4 to 40 rpm over 5.5 min). The latency to fall from the rod was averaged over 5 trials in individual mice.

#### Responses to noxious stimuli on the hindpaw

Each mouse was acclimatized in the test apparatus for 1-2 hours before the application of stimuli.

#### Mechanical stimuli

A series of calibrated von Frey filaments was used to apply mechanical stimuli to the plantar surface of the hindpaw. We used the up-down paradigm to determine the 50% withdrawal threshold (Chaplan et al., 1994).

#### Thermal and cold stimuli

We recorded the latency to lick the hindpaw and/or to jump on a 55°C hot plate. The cut-off time was 20 s. We also measured the paw withdrawal latencies to radiant heat stimuli applied to the plantar surface of the hindpaw (Hargreaves et al., 1988). The cold plantar assay was used to compare the response latencies of WT and TRESK KO mice to cold stimuli on the plantar surface of the hindpaw (Brenner et al., 2012).

#### Chemical stimuli

Mice were injected with 0.3 µg capsaicin (in 10 µl saline with 1% DMSO) in the plantar surface of one hindpaw. We quantified the time spent licking the treated paw within 5 min after the injection.

#### Visceral pain test

We counted the number of abdominal stretches that occurred within 30 min after i.p. injection of 5.0 ml/kg 0.6% acetic acid, a stimulus that produces visceral pain with inflammation (Cao et al., 1998).

### Responses to facial noxious stimuli

#### Eye blink responses to thermal stimuli

Mice were well-habituated to the test apparatus and extensively handled by the experimenter for a week before testing. On the test day, the experimenter gently held the mouse on the palm and delivered the thermal stimulus by blowing air to one eye at 0.5 liter/min for 10 s. The number of eye blinks was recorded by a video camera and quantified off-line. The distance between the air outlet (6.35 mm diameter) and the eye was 2-3 mm. The air was heated to various temperatures (20-55°C) by passing through plastic tubing submerged in heated water bath. The air temperature was measured by a thermal probe at the outlet. Mice were tested every 3 days. Each eye was tested twice per day, first with RT air and 1-2 hour later with air at a higher temperature. Results from left and right eyes were averaged for individual mice.

#### Cheek acetone test

We measured the acute nocifensive responses to acetone-evoked evaporative cooling on mouse cheek (Constandil et al., 2012; Trevisan et al., 2016). Mice were habituated individually in clear plexiglass boxes (11 × 11 × 15 cm) situated in front of three angled mirrors for at least 1 hour for 3 consecutive days and were extensively handled by the experimenter. The day before testing, both cheeks were shaved (6.5 × 12 mm) under brief anesthesia (2% isoflurane in 100% oxygen). On the test day, mice were habituated for 1 hour before the assay. We applied acetone (15 µl) to the shaved cheeks and immediately returned mice to the boxes. Time spent wiping the treated area was recorded by a video camera and quantified off-line. For individual mice, acetone was applied alternatingly to both cheeks at > 10 min interval, and the duration of behavior was averaged from 4 applications.

#### Cheek capsaicin injection

Mice were acclimated and shaved as in the facial acetone test. On the test day, we habituated mice in the clean boxes for 1 hour and then injected 1 µg capsaicin (in 20 µl saline with 1% DMSO) or vehicle intradermally in one cheek (Akiyama et al., 2010; Shimada & LaMotte, 2008). Mice were immediately returned to the boxes and recorded by video cameras for 40 min. Total numbers of forepaw wiping of the injected cheek were quantified off-line.

#### Operant behavioral responses to thermal stimuli

Responses to facial thermal stimuli were tested in the OPAD (Stoelting), which uses a reward-conflict paradigm that allows a mouse to choose between receiving a reward (sweetened condensed milk diluted 1:2 with water) and escaping an aversive stimulus (50°C thermal stimuli on the cheeks), thereby controlling the amount of pain it feels. First, mice underwent 4 training sessions in two weeks. The day before each session, mice were food deprived for 16 hours and both cheeks were shaved. The next day, individual mice were trained in the OPAD for 20 min to learn to voluntarily press the cheeks onto the Peltier bars set at 33°C in order to drink sweetened milk (Anderson et al., 2013). Mice that consistently licked more than 600 times / per session were used to test facial thermal responses. Trained mice underwent 2 test sessions 3 days apart. During each test session, the thermode temperature varied every 3 min as follows: 33°C–50°C– 33°C–50°C–33°C. The number of licks and the number of thermode contacts at each temperature were recorded and averaged in individual mice.

#### Withdrawal responses to facial mechanical stimuli

Mice were well-habituated to the test room and extensively handled by the experimenter for at least two weeks before testing. The hair on the forehead (above and between two eyes) was shaved the day before testing. On the test day, the experimenter gently held the mouse on the palm with minimal restraint and applied the calibrated von Frey filament perpendicularly to the shaved skin, causing the filament to bend for 5 seconds. A positive response was determined by the following criteria as previously described: mouse vigorously stroked its face with the forepaw, head withdrawal from the stimulus, or head shaking (Elliott et al., 2012). We used the up-down paradigm to determine the 50% withdrawal threshold (Chaplan et al., 1994). The responses of female WT and KO mice to von Frey filaments were measured before and 0.5-1 hour after the i.p. injection of TRPV1 antagonist AMG9810 (Sigma, 10 mg/kg in 1% DMSO). This dose of AMG9810 effectively inhibited cheek wiping induced by intradermal injection of 1 µg capsaicin (data not shown).

#### Mouse model of episodic migraine

Craniectomy was performed on 9-15 weeks old WT and TRESK KO mice as described in a previous study (Huang et al., 2016) and in the section of retrogradely labeling of dural afferent neurons above. We applied 20 µl of sterile saline to the dura and let mice recover from surgery for 7 days. Mice were housed individually in the animal facility and were transported to the testing room for acclimation and handling every day. On the test day, mice in their home cages were habituated in the testing room for 1 hour and were briefly anesthetized with isoflurane. The polypropylene cap was removed without disturbing the skin suture and 20 µl of vehicle (saline with 2% DMSO) or IScap solution (see below) was applied onto the exposed dura. Subsequently, a new sterile polypropylene cap was secured over the ring with bone wax to cover the dura. After recovery from anesthesia, mice were returned to the home cages placed in front of a 3-way mirror to ensure that the head-directed behavior be recorded at all body positions. Home cage behaviors were recorded 0.5-2.5 hours after the dural application. Mice were recorded one at a time in the absence of the experimenter and other mice. Digital video files were quantified off-line by experimenters blinded to the genotypes of mice or the treatments that mice received. The entire 2-hour video was watched and scored. The total number and the duration of forepaw wiping and hindpaw scratching within the mouse V_1_ dermatome (including the scalp and periorbital area) were quantified.

The IScap solution contained 0.5 mM capsaicin, 1 mM bradykinin, 1 mM histamine, 1 mM serotonin (5-HT), and 0.1 mM prostaglandin E2 (PGE_2_) in saline with 2% DMSO. All chemicals were purchased from Sigma, dissolved in H_2_O (bradykinin, histamine, and 5-HT) or DMSO (capsaicin and PGE_2_) at 100x concentrations and stored at −80°C in aliquots. IScap solution was freshly prepared from the stock solutions on the test day.

### Statistical analysis

For *in vitro* experiments, data from each group contained neurons from at least 3 independent cultures using at least 2 mice each time. For behavioral experiments, power analysis is conducted to estimate sample size with > 80% power to reach a significance level of 0.05. For IHC experiments, sample sizes were estimated based on our prior experience.

All data are reported as mean ± standard error of the mean. The Shapiro–Wilk test was used to check data normality. Statistical significance between experimental groups was assessed by Fisher’s exact test, χ^2^ test, two-tailed t-test, analysis of variance (ANOVA) (1-way or 2-way, with or without repeated-measures [RM]) with the post hoc Bonferroni test where appropriate, using Origin and Statistica (from OriginLab and StatSoft, respectively). The non-parametric Mann–Whitney U test or the Kruskal-Wallis ANOVA was used where appropriate to analyze the differences in the soma size distribution and the withdrawal threshold to mechanical stimuli. Differences with *p* < 0.05 were considered statistically significant.

## Acknowledgements

The authors thank Dr. Daizong Li for the generation of the TRESK antibody; Drs. Hongzhen Hu, Jing Feng, Jin Hua, Ms. Megan E Cloud, Katherine Tsay, and Alicia Liu for technical assistance; as well as Drs. Christopher J Lingle and Joe Henry Steinbach for valuable comments on the manuscript. We also thank the Alafi Neuroimaging laboratory and the Pain Center Animal Behavior Core at Washington University. This work was supported by National Institutes of Health grants NS083698, NS087321 and NS103350 (to YQC), and a Xiangya Scholarship Fund (to XHJ).

## Author Contributions

Conceptualization, Y.Q.C. and Z.H.G.; Methodology, Z.H.G., C.S.Q., J.T.Z, F.X.L., and Q.L.; Investigation, Z.H.G., C.S.Q., J.X.H. and J.T.Z; Resources, A.D.; Writing – Original Draft, Y.Q.C. and Z.H.G.; Writing – Review & Editing, Y.Q.C., Z.H.G., Q.L., and A.D.; Supervision, Y.Q.C. and Z.H.G.; Funding Acquisition, Y.Q.C.

